# Meta-analysis of the effects of clinically-effective therapeutics in the preclinical migraine model as a tool for design optimisation

**DOI:** 10.1101/2022.07.05.498780

**Authors:** Antonina Dolgorukova, Ekaterina Protsenko, Julia Isaeva, Victoria Gagloeva, Elena Verbitskaya, Alexey Y. Sokolov

**Affiliations:** Valdman Institute of Pharmacology, Pavlov First Saint Petersburg State Medical University, Saint Petersburg, Russia; Russian Medical Academy of Continuous Professional Education, Moscow, Russia; Laboratory of Cortico-Visceral Physiology, Pavlov Institute of Physiology of the Russian Academy of Sciences, Saint Petersburg, Russia

**Keywords:** Electrophysiology, Migraine, Trigeminovascular system, Neuronal activity, Animal model, Sample size calculation, Systematic review, Meta-analysis

## Abstract

The low reliability of the preclinical study’s findings is of critical concern. The possible sources include poor experimental design and a lack of measures to reduce the risk of bias. In this study, we focused on anti-migraine drug discovery and a particular animal model with the aim to contribute to the elimination of these sources in future research. We performed a systematic search of controlled studies testing established migraine treatments in the model of trigeminovascular nociception (EMTVN) and meta-analysis for the main outcomes to estimate the overall effect sizes. In 13 studies reporting on 21 experiments, anti-migraine drugs significantly decreased trigeminovascular nociceptive traffic compared with a control intervention. Considering these effects biologically relevant, we used them in sample size calculation for future experiments. To refine the EMTVN and inform its users, we explored the impact of methodological features on the outcome and revealed several factors potentially impacting the results obtained in this model. We also assessed the internal validity of the included studies and found that the selection bias, particularly, the lack of randomisation, is likely a main source of bias. Based on our findings, we discuss the translational potential of the EMTVN and suggest what should be addressed for its improvement. We believe that this work highlights the importance of systematic reviews and meta-analyses as tools for design optimisation in animal research.

## 1. Introduction

Animal models serve as the backbone of drug research and development, but there is a growing concern about the low reliability and reproducibility of preclinical study’s findings (Button et al., 2013; Macleod and Mohan, 2019). The possible causes include biased study design, data collection, and analysis as well as poor reporting, and insufficient sample size, which results from the lack of prior calculations. The latter is of particular importance, since the probability that a finding of an underpowered study is false is much greater than that of a study with conventional power (Button et al., 2013). The main difficulty in calculating the sample size is that it usually requires specification of the effect size to be detected. It ideally should represent a meaningful difference between the treatment and control, but this is often unclear how large this difference must be to be considered meaningful. In this review, we have hypothesised that for a certain animal model, the summary effect of previously tested clinically effective drugs may be considered biologically relevant and serve as a reference for sample size calculation. To examine how feasible this is, we have chosen the field of anti-migraine drug discovery research and one particular technique used in our laboratory – the electrophysiological model of trigeminovascular nociception (EMTVN).

Mechanisms of migraine involve complex interplay between various brain regions, and the exact chain of events is still unknown, complicating preclinical modeling of this disorder (Dodick, 2018; Harriott et al., 2019). However, the key process in its pathogenesis has been established: it is the activation of the trigeminovascular system (TVS) (Ashina et al., 2019). The TVS consists of peripheral and central trigeminal neurons functionally integrated with cranial blood vessels and plays a major role in the integration of pain signals and their transmission to higher brain centres (Ashina et al., 2019; Dodick, 2018). The EMTVN recapitulates TVS activation in the whole live animal by stimulation of the meninges richly innervated by trigeminal afferents with extracellular recording from the responsive first-, second-, or third-order trigeminovascular neurons in the trigeminal ganglion, trigeminocervical complex, and thalamic nuclei, respectively in real time (Harriott et al., 2019). The assessment of the stimulation-evoked and ongoing neuronal activity in the EMTVN allows direct examination of drug effects on the transmission of pain signals along the trigeminovascular pathway.

In this study, we have performed a systematic search of previously published controlled studies reporting testing of clinically established anti-migraine medications in the EMTVN. We have estimated the summary effects of these drugs with meta-analysis and have calculated sample sizes that allow to detect such effects in future experiments. To refine the model and inform its users, we have explored the impact of methodological features on the outcome, as well as the possible sources of bias and suggest several steps needed for study design optimisation.

## 2. Materials and methods

The full protocol of this study with prespecified methods is available at PROSPERO website (CRD42021276448; https://www.crd.york.ac.uk/prospero/display_record.php?ID=CRD42021276448). The amendments include network analysis, performed to examine whether there are clusters formed by the study’s authors, and an addition to the sample size calculation, providing sample sizes required under multiple conditions. The detailed description and reasons for each amendment can be found in the respective section below.

### 2.1. Search

In September 2021, we systematically searched three online databases (PubMed, Scopus, and eLibrary) with no language restrictions to identify publications reporting testing of clinically established anti-migraine medications in the EMTVN. We updated the search in December 2021 at the end of the data extraction phase, as was planned in the study protocol. The search strategy was based on terms relevant to the model of interest, which are detailed in the Supp. 1. Search results were limited to publication date from 1986, as in this year two research groups have shown for the first time the existence of trigeminal neurons responsive to electrical stimulation of intracranial blood vessels (Davis and Dostrovsky, 1986; Strassman et al., 1986), the basis for the future emergence of the EMTVN. The records found were imported into the reference management software (Endnote X7). Duplicate references were removed using either the Automated Systematic Search Deduplication tool developed by the CAMARADES group (Hair, 2019; Hair et al., 2021) (for the records identified during the initial search) or manually (for the records identified during the updated search).

### 2.2. Inclusion and exclusion criteria

We included original peer-reviewed full-length research papers reporting experiments performed using the EMTVN, which was defined as follows: neurons or neuronal clusters responsive to dural stimulation (mechanical, thermal, or electrical) were extracellularly recorded in the trigeminal ganglion, trigeminocervical complex, or thalamic nuclei in whole live animals under general anaesthesia. There were no restrictions on the methodological features eligible for inclusion. The eligible drugs were acute pharmacological interventions recommended for clinical use in migraine according to national guidelines and/or headache societies: European: (Steiner et al., 2019); US: American Headache Society Position Statement (2019); UK: National Headache Management System for Adults (BASH, 2019); and Russian: (Filatova et al., 2020) and/or approved for clinical use in the US, UK, Europe or Russia that were tested with any systemic route of administration. The list of drugs that meet these criteria is available in the study protocol. We excluded combinations of two or more interventions irrespective of the rationale (e.g., pharmacological antagonism or additivity/synergism, pretreatment with sensitising agents). The eligible animals were laboratory rodents of any strain, age or sex. We excluded experiments performed in genetically modified organisms, not experimentally naive animals, animals with lesion to any CNS area or structure. The eligible outcomes were changes from baseline of the ongoing neuronal activity and neuronal responses evoked by dural stimulation after drug and control administration. We excluded experiments performed without appropriate control (e.g., historical controls) or with mixed designs.

### 2.3. Study selection

Each record was screened by two independent reviewers. First, titles and abstracts identified through searching were screened, and ineligible studies (e.g. reviews, abstracts, comments, books, etc.) and those clearly made without the EMTVN were excluded. For the remaining reports we obtained full paper copies. Publications not reporting at least one experiment with eligible model, intervention, and animals were excluded. The experiments from the remaining records were assessed for eligible control and outcome measures. With respect to all screening stages, any discrepancies between reviewers were resolved through discussion, or by consulting a third investigator.

### 2.4. Risk of bias and quality assessment

Two independent reviewers assessed the risk of bias of the included studies in accordance with the SYRCLE Risk of Bias tool with some adaptations by recording the reporting of randomisation; sample size calculation; the absence of between-group difference in the outcome at baseline and/or accounting for baseline fluctuations in the analysis; blinded assessment of outcome; the information for attrition, reporting, and design-specific bias assessment; the information for influence of funders assessment. Risk was scored high, low, or unclear based upon the criteria presented in Table 1. Reporting of compliance with animal welfare regulation and the approval of the study protocol by an institutional animal care and use committee (or equivalent) were also collected. Discrepancies were reconciled by a third reviewer.

**Table 1.** Risk of Bias assessment.

| **Bias** | **Low** | **High** | **Unclear** |
| --- | --- | --- | --- |
| Selection bias: randomisation | Valid method reported | Inappropriate method reported | Randomisation not reported; method not reported |
| Selection bias: sample size calculation | Valid method reported | Inappropriate method reported | Sample size calculation not reported; method not reported |
| Selection bias: similarity at baseline and/or accounting for baseline fluctuations in the analysis | The absence of between-group difference in the outcome at baseline or accounting for baseline fluctuations in the analysis reported | Between-group difference reported without accounting in the analysis | Between-group comparisons of baseline data and accounting for baseline fluctuations in the analysis not reported |
| Performance bias: blinding | Blinded assessment of outcome reported | Reported that blinding was not performed | Blinded assessment of outcome not reported |
| Attrition bias: incomplete outcome data | Number of units tested is equal to the number of units for which the outcome is reported | Number of units tested is not equal to the number of units for which the outcome is reported; animal/neuron exclusions reported without providing a reason; exclusion of non-responders reported | Not enough data to score. |
| Reporting bias: selective outcome reporting | Outcomes reported in methods and results section of the article are similar | Outcomes reported in methods and results section differ | Not enough data to score. |
| Influence of funders | The authors declare that there was no conflict of interests | One or more authors are affiliated with the organization that provided the studied drug; the research was supported by organization that provided the studied drug | Not enough data to score. |
| Design-specific bias | Parallel study design or within-subject designs, which have been adequately addressed in the analysis and the absence of biological replicates reported | Biological replicates or within-subject designs, which have not been adequately addressed in the analysis reported | Not enough data to score. |

*By biological replicates is meant the recording of more than one neuron/neuronal cluster in one animal.*

### 2.5. Data extraction and reconciliation

Details of each publication provided in Table 2 were extracted by two independent reviewers into SyRF (syrf.org.uk). In the case outcome data were presented graphically, we used Adobe measuring tool to determine values. When multiple time points were presented, the time point that showed the greatest difference between the control group and treatment group was extracted. If a drug was studied at multiple doses, we extracted the effects of the maximal dose. When the type of variance was not reported, it was characterised as SEM. To obtain additional information on poorly reported items we contacted the corresponding author.

**Table 2.**
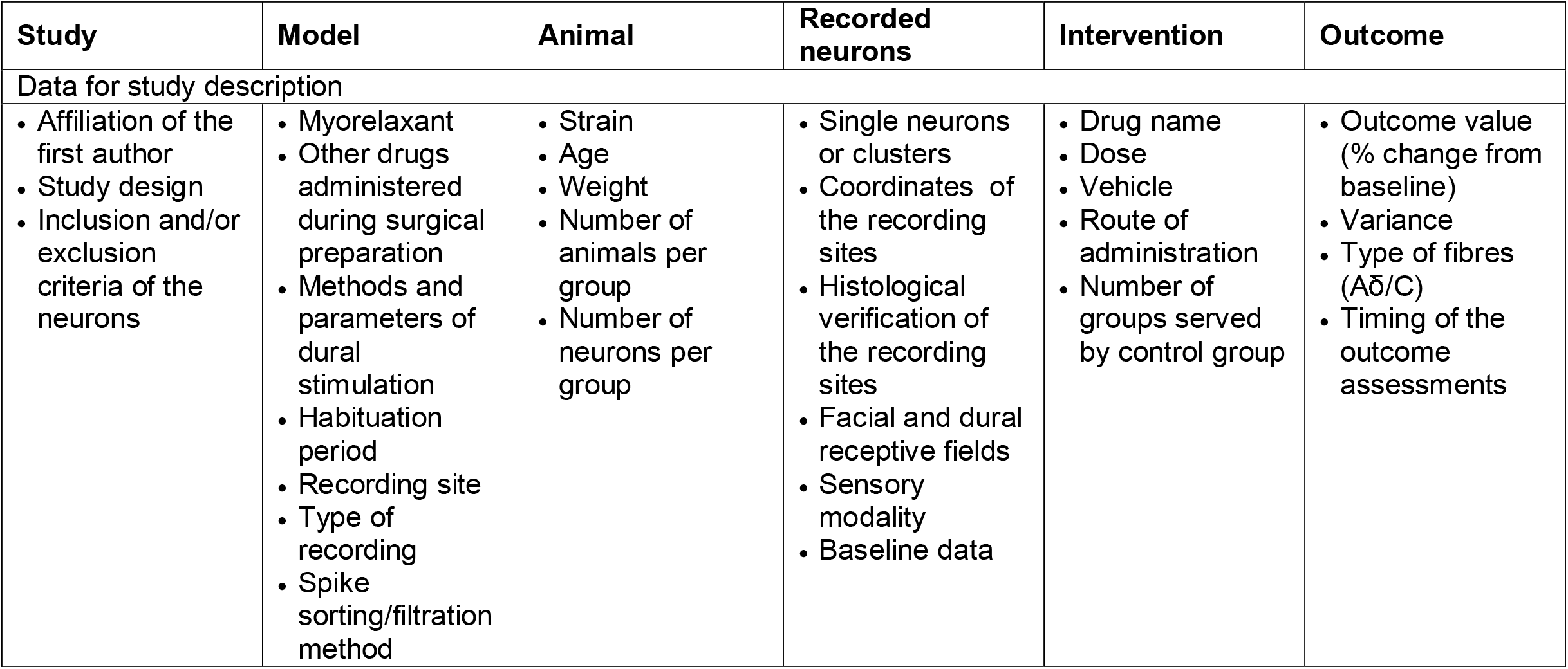
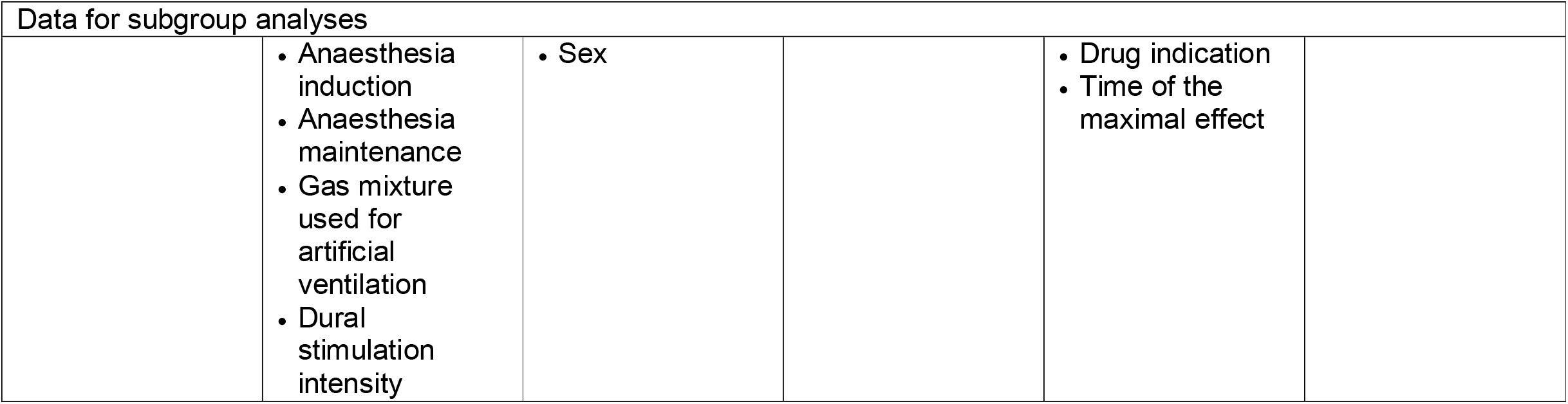
Data extracted from each included publication.

| Study | Model | Animal | Recorded neurons | Intervention | Outcome |
| --- | --- | --- | --- | --- | --- |
| Data for study description |  |  |  |  |  |
| Affiliation of the first author<br>Study design<br>Inclusion and/or exclusion criteria of the neurons | Myorelaxant<br>Other drugs administered during surgical preparation<br>Methods and parameters of dural stimulation<br>Habituation period<br>Recording site<br>Type of recording<br>Spike sorting/filtration method | Strain<br>Age<br>Weight<br>Number of animals per group<br>Number of neurons per group | Single neurons or clusters<br>Coordinates of the recording sites<br>Histological verification of the recording sites<br>Facial and dural receptive fields<br>Sensory modality<br>Baseline data | Drug name<br>Dose<br>Vehicle<br>Route of administration<br>Number of groups served by control group | Outcome value (% change from baseline)<br>Variance<br>Type of fibres (Aδ/C)<br>Timing of the outcome assessments |

| Data for subgroup analyses |  |  |  |  |
| --- | --- | --- | --- | --- |
|  | • Anaesthesia induction<br>• Anaesthesia maintenance<br>• Gas mixture used for artificial ventilation<br>• Dural stimulation intensity | • Sex |  | • Drug indication<br>• Time of the maximal effect |

We compared the data extracted by two reviewers, and a third independent reviewer reconciled discrepancies. For outcome data extracted from graphs, if the values extracted by the two reviewers differed by >10%, they required reconciliation, otherwise, we calculated the means of the two values.

### 2.6. Data analysis

The data were separated by outcome into two data sets: the first with experiments where responses evoked by dural stimulation were assessed, and the second with experiments where ongoing neuronal activity was measured. We used an individual experiment (defined as a drug tested using a particular route of administration on an animal/neuronal cohort) as an observational unit in the analyses. The absolute difference in means (MD) was used as an effect size measure. The standard errors of the effect sizes were calculated using formulas 5 and 6 in Vesterinen et al. (2014). If different measures of the neuronal activity were reported (e.g. for Aδ- and C-fibres), we combined the data using the formulas 24 and 25 in Vesterinen et al. (2014) to provide a single effect per experiment and avoid artificial inflation of the numbers of units and within-unit correlation among the effect sizes. Since both data sets included publications reporting more than one experiment per study with a shared control group, i.e. with dependent effect size estimates, we fitted a three-level meta-analytic model with experiments nested within studies, resulting in a mixed-effects model with a random effect corresponding to a study and a random effect corresponding to an experiment within a study, and computed robust variance estimates (Hedges et al., 2010) using metaphor package in R (Viechtbauer, 2010). For graphical presentation and results description we calculated 95% confidence intervals (95%CI).

For each data set we calculated the percentage of variability that can be attributed to the overall heterogeneity (which is the sum of between- and within-study heterogeneity) rather than a chance (sample variance) using Higgins’s I^2^ (0–25% was interpreted as very low heterogeneity; 25–50% as low heterogeneity; 50–75% as moderate heterogeneity; and >75% as high heterogeneity) and examined the distribution of variance over experiment level and study level (Viechtbauer, 2022). To refine the modelling of EMTVN and inform future studies, we assessed the extent to which methodological features (Table 2) and the reporting of measures to reduce bias explain the observed heterogeneity by subgroup analyses (meta-regression). To test the robustness of the overall results, that is to ensure that there are no outlying experiments that may have an excessive influence on the overall effect estimates, we performed sensitivity analyses. For the sake of reliability and robustness, these exploratory analyses were performed only for data sets with ≥10 experiments and high heterogeneity. According to our results, the observed heterogeneity could not be explained by any of the pre-specified factors, and besides, during this work, we noticed that there may be few distinct research groups. To examine whether there are clusters formed by the study’s authors, we conducted and visualised a network analysis.

We detected outliers using standardized (deleted) residuals and influential cases based on Cook’s distances. The advantage of the deleted residuals is that they are more sensitive to detecting outliers (Schmid et al., 2020). We assumed that residuals larger than ±1.96 indicate experiments that does not fit the assumed model, i.e. represent outliers, and Cook’s Distance over 4/n (where n is the total number of data points) indicates influential cases. We removed influential outliers from the data set and rerun the analyses. All analyses were performed using R programming language (Harrer et al., 2021; Schmid et al., 2020; Viechtbauer, 2010).

### 2.7. Sample size estimation for future studies

For each outcome, we calculated an overall effect size, dividing the estimated mean difference between the treatment and control by the average pooled standard deviation (SD) of that outcome. This value we used as an input to the standard sample-sizing formula for the Wilcoxon-Mann-Whitney test (based on the two-sided alpha = 0.05 and 80-100% power, allocation ratio 1:1) to obtain the number of animals required in future experiments. Our analysis, however, indicated that the pooled SD can considerably vary from one laboratory to another, while the overall effect size is likely to be overestimated. Consequently, we assumed that it would be more useful and informative to calculate sample sizes required under various conditions, and made an addition to the sample size calculation pre-specified in the protocol. For this, we ranked the pooled SDs from high to low using 80^th^, 50^th^ and 20^th^ rank percentiles (Currie et al., 2019) and calculated the sample size allowing to detect 80% and 50% of the overall effect given the estimated outcome variance.

### 2.8. Publication bias

We assessed the potential publication bias by estimating the asymmetry of funnel plots of the experiment level data using visual inspection and Egger’s regression test if there are sufficient numbers of comparisons (n ≥ 20). For this, we fitted a multi-level meta-regression model with a measure of effect size precision (standard error) as a predictor and used robust variance estimates to handle dependence (Rodgers and Pustejovsky, 2020).

## 3. Results

### 3.1. Identification of publications

Our initial systematic search (September 2021) identified 6127 publications, and the updated search (December 2021) identified the additional 44 articles (Figure 1). After the removal of duplicate references and initial screening based on titles and abstracts, full-texts of 431 records were assessed for eligibility. Of them, we included 18 records (17 studies and one PhD thesis) reporting on 28 experiments that met the inclusion criteria.

**Figure 1.**
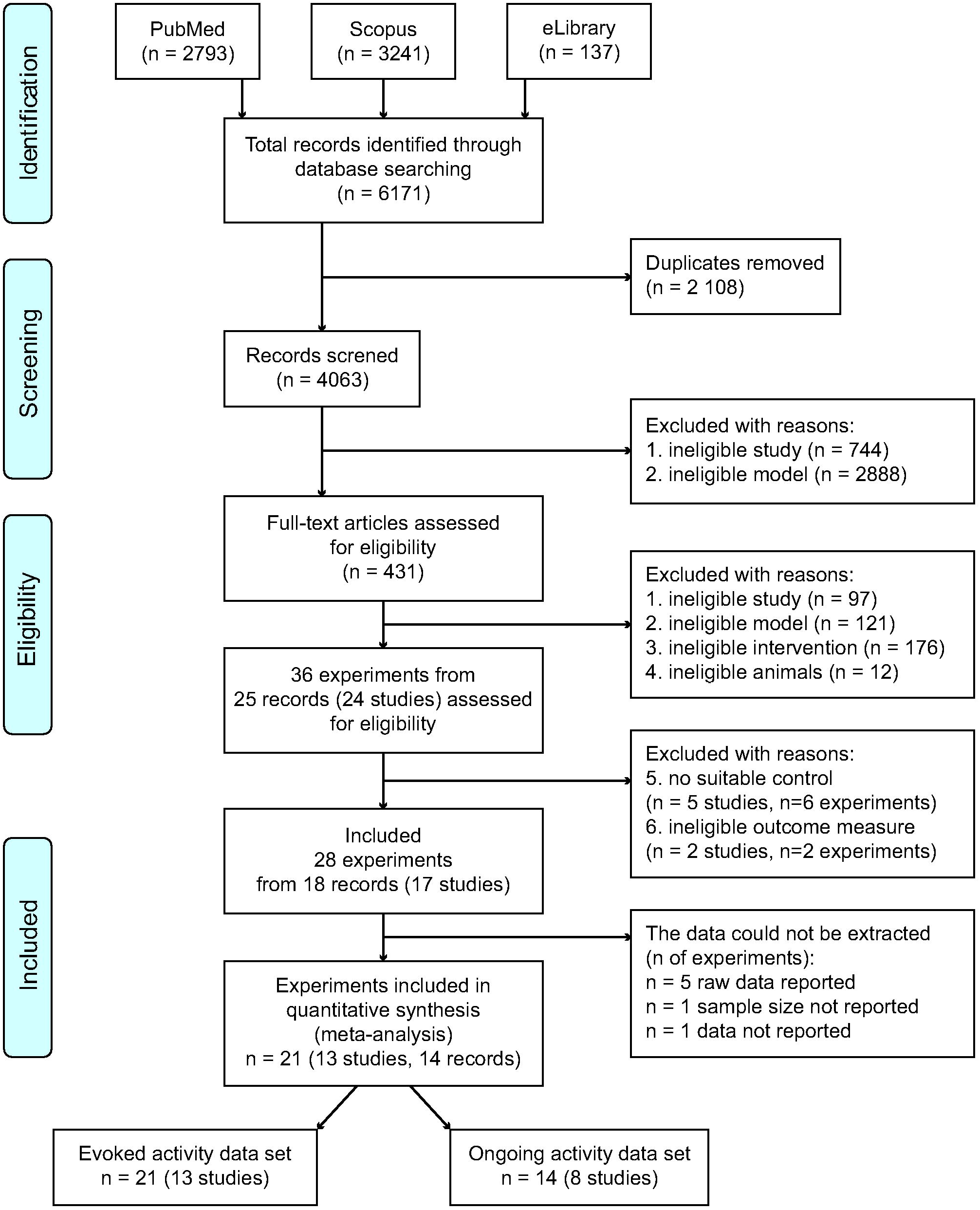
A flow diagram of studies identification and screening process reported in accordance with the PRISMA guidelines. The search was performed in 3 electronic databases: PubMed, Scopus, and eLibrary up to December 14, 2021. The diagram provides the breakdown of records through deduplication, screening, and eligibility until final inclusion in quantitative synthesis.

### 3.2. Missing information

We could not extract data from 7 experiments reported in 4 studies. Melo-Carrillo et al. (2017) and Burstein et al. (2005) reported raw data, Charbit et al. (2009) did not report the number of units in the control group, and Shields and Goadsby (2006) did not report the effect of the control intervention numerically.

In 6 of the remaining 13 studies reporting on 21 eligible experiments, some data (values and/or variance and/or n in groups) for one or both outcomes were missing. In one study (Akerman and Romero-Reyes, 2019), the effects of the drugs on the evoked activity were reported in absolute values, and the authors kindly provided the data expressed as % changes from baseline as well as numbers of units with C-fibre inputs. Dolgorukova et al. (2020) reported medians and interquartile ranges, and means with SDs were obtained from the authors. The corresponding authors of the remaining 4 studies did not respond to our data request. Of them, Bergerot et al. (2007) did not report the effects of the drugs tested on the ongoing activity numerically (it was not affected); Oliveira et al. (2016) did not report the variance and the effect of the control intervention on the ongoing activity (it was reduced by the drug maximally by 58% of baseline); Andreou and Goadsby (2011) did not report the number of units with C-fibre inputs; Zhao et al. (2018) did not report the effects of the drugs tested on the C-fibre responses numerically (they were not affected).

Overall, at least one measure of the dural stimulation-evoked (DS-evoked) activity was reported in all 21 experiments, and the ongoing activity was measured in 16 experiments; however, we could not extract data from two of them (Bergerot et al., 2007; Oliveira et al., 2016). The meta-analysis thus included 13 studies with 21 eligible experiments assessing the effects of anti-migraine drugs on the DS-evoked neuronal activity and 14 experiments reporting on drugs’ effects on the ongoing firing rate.

Regarding the pre-specified factors for subgroup analyses, authors of three studies supplemented the description of dural electrical stimulation (Summ et al. (2021), Akerman and Romero-Reyes (2019), and Vila-Pueyo et al. (2021) used threshold intensity); one author from our group also provided several poorly reported items for two studies (Sokolov et al. (2008) used urethane and alpha-chloralose for anaesthesia maintenance; Sokolov et al. (2013) used room air for artificial ventilation).

### 3.3. Study characteristics

According to the first author’s affiliations, the included 13 studies were performed in 4 different countries (Supp. 2, Table 1). All the experiments (n = 21) were made in male Sprague-Dawley (n = 15) or Wistar (n = 6) rats of 220 - 440 g.

The studies used 6 different anaesthesia regimens: inhalation anaesthetics halothane and isoflurane, sodium pentobarbital, and urethane with or without alpha-chloralose were used for anaesthesia induction; propofol, sodium pentobarbital, and urethane with or without alpha-chloralose were used for anaesthesia maintenance (Figure 2, A-B). Artificial ventilation was performed in all studies, but one of them (Cumberbatch et al., 1997) did not specify the gas mixture used (Figure 2, C). Myorelaxation was reported in 8/13 studies. One of the 13 studies (Farkas et al., 2015) reported other drugs administered during surgical preparation (atropine, 50 ug/kg, s.c.; acetazolamide, 10 mg/kg, i.p.; lignocaine, 10% w/w, routinely on skin incisions, in the region of femoral artery cannulation, and the ear canal before mounting in the stereotaxic apparatus). The habituation period (period of time during which animals rested after surgical preparation) was reported in only 3/13 studies and was 30 or 60 min.

**Figure 2.**
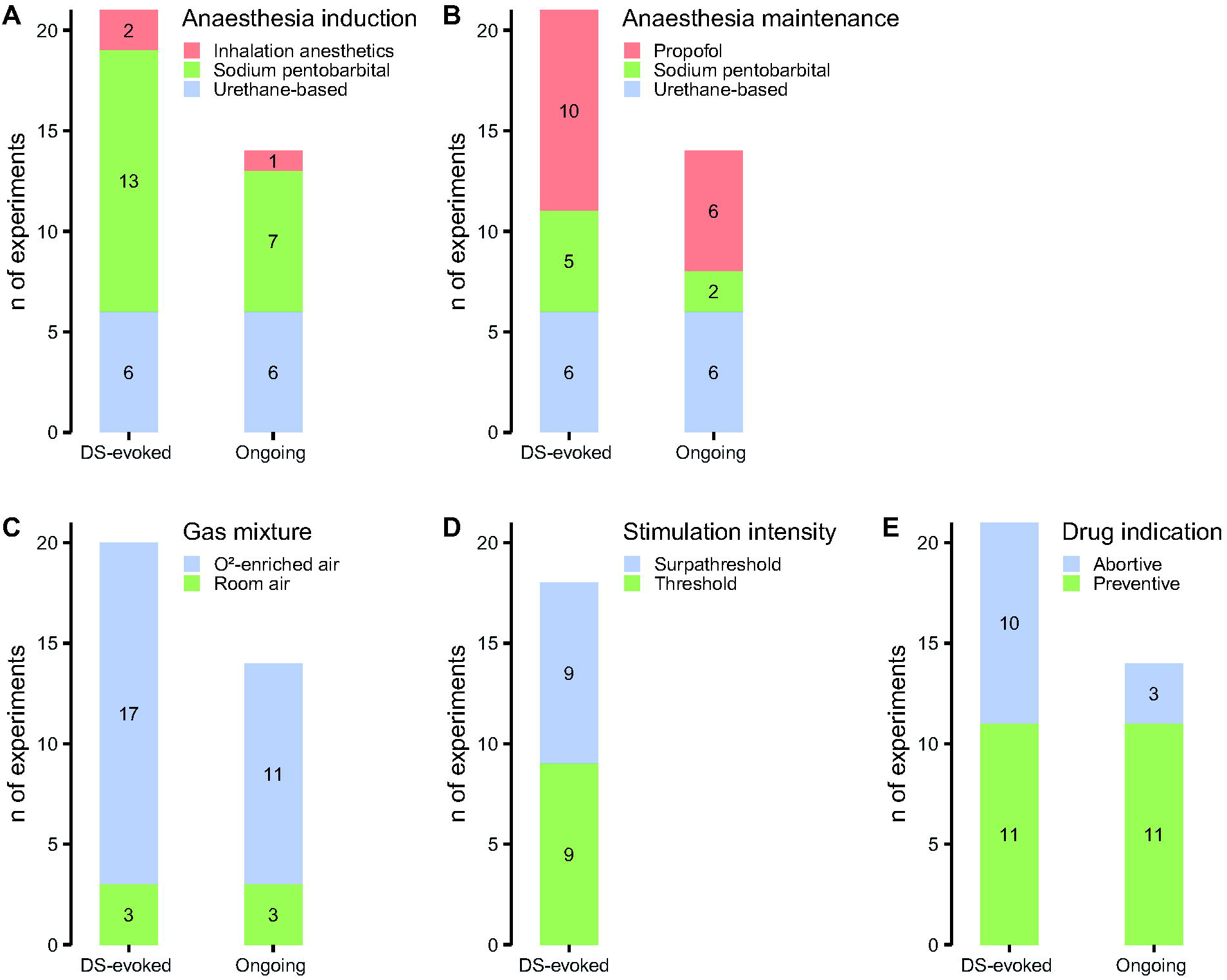
Categorical factors assessed in the subgroup analyses. Anaesthesia induction (A) and maintenance (B), gas mixture used for artificial ventilation (C), intensity of electrical dural stimulation (D), and the indication of studied drugs (E). Urethane-based anaesthesia means that urethane was used with or without alpha-chloralose.

For the assessment of DS-evoked activity of trigeminovascular neurons, all studies used electrical stimulation of the meninges at threshold (4 studies, 9 experiments) or suprathreshold (7 studies, 9 experiments) intensity (Figure 2, D). The intensity of electrical pulses was not stated in 2/13 studies (Bergerot et al., 2007; Hoffmann et al., 2019) reporting on three eligible experiments.

Overall, 290 neurons or neuronal clusters were recorded in the trigeminocervical complex (12 studies, 19 experiments) or ventroposteromedial thalamic nucleus (2 studies, 2 experiments). Eight of 13 studies reported the coordinates of recording sites, but only 3 of them provided complete information on spatial relationships (anterior-posterior, medial-lateral, and dorsal-ventral coordinates). Histological verification of the recording sites was performed in 4 studies, including these with recordings in the ventroposteromedial thalamic nucleus. Three studies (Andreou and Goadsby, 2011; Bergerot et al., 2007; Oliveira et al., 2016) reported biological replicates (i.e. when more than one neuron/cluster was recorded per animal), but none of them clarified whether the units were recorded simultaneously or sequentially.

Inclusion and/or exclusion criteria of neurons (other than sensitivity to dural stimulation) were reported in 11/13 studies. Predominantly, these were the requirements for input from the facial skin (Supp. 2, Table 2). The characteristics of the studied neuronal populations are presented in Table 3. It should be noted that most studies included other experiments, which were not eligible for this review. In these cases, the characteristics of the neurons were reported for all studied units.

**Table 3.**
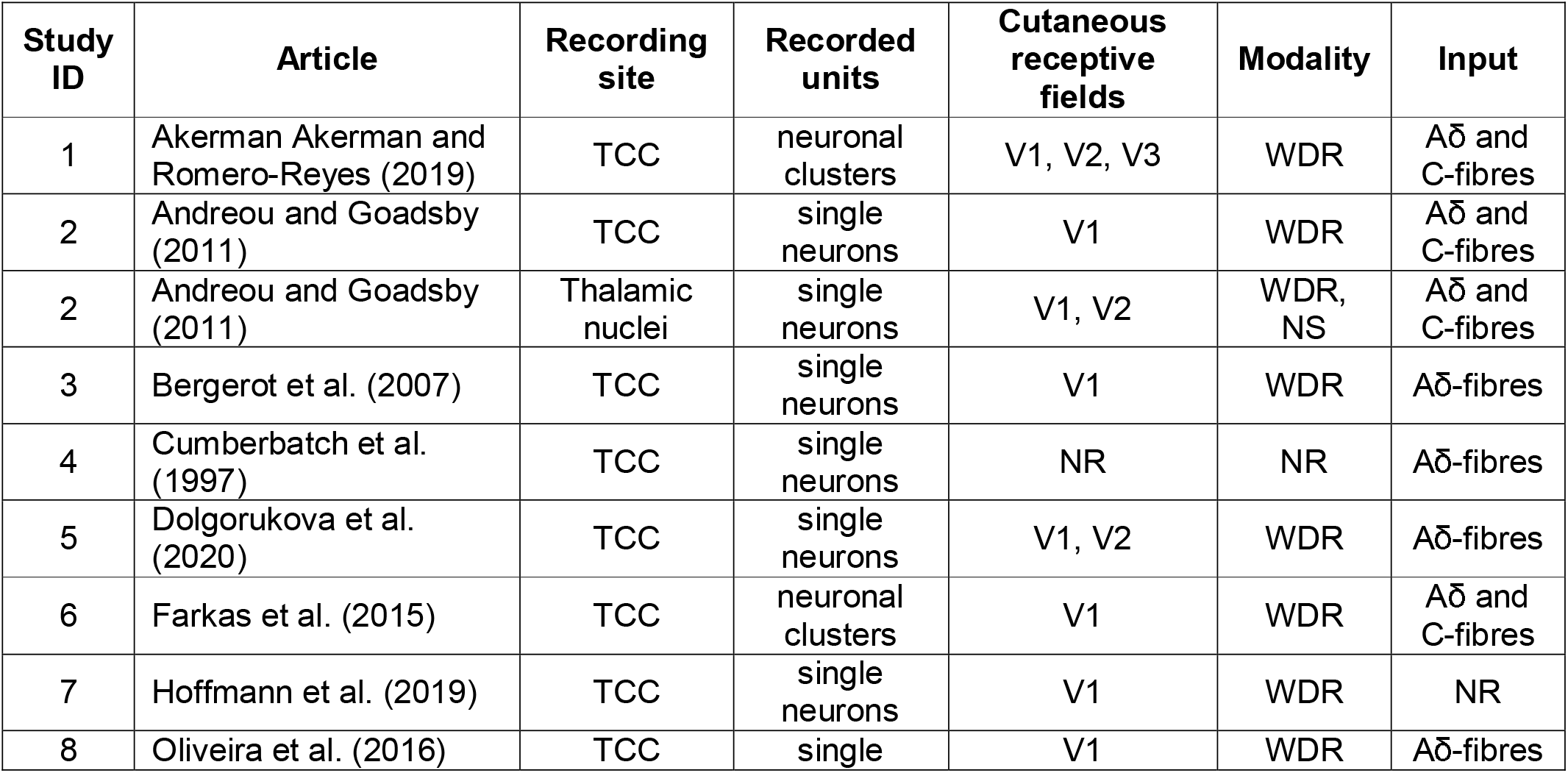
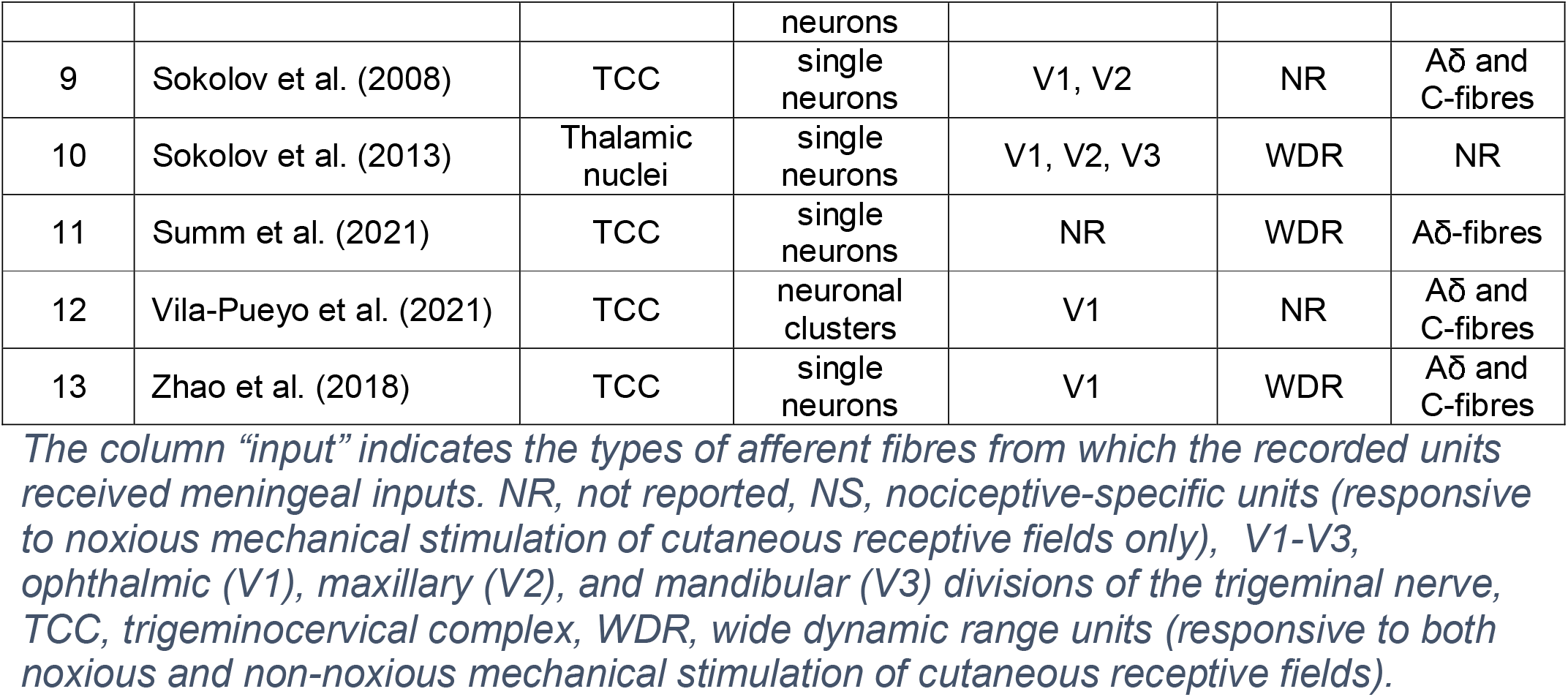
Characteristics the studied neuronal populations.

Five of 13 studies reported average baseline values for the ongoing activity of all examined neurons, and 4/13 studies reported average baseline values for the evoked activity (Supp. 2, Table 13-14).

The included 13 publications assessed the effects of 13 abortive and preventive anti-migraine drugs (**Error! Reference source not found**., E). The most frequently tested drugs were naratriptan, propranolol, and valproate (each tested in 3 experiments). The time of the maximal effect varied between 5 - 90 min after the drug administration. The first outcome assessment was performed from 0 to 15 min after treatment, and the last was made 30 to 120 min after treatment.

In Supp. 2 we provide a detailed description of the included studies and experiments.

### 3.4. Meta-analysis

The meta-analysis was made for each of the two data sets: the first included 13 studies with 21 eligible experiments assessing the effects of anti-migraine drugs on the DS-evoked neuronal activity and the second included 8 studies with 14 eligible experiments reporting on drugs’ effects on the ongoing firing rate (Supp. 3, Table 1-2). Administration of established anti-migraine therapeutics led to a significant reduction of both the DS-evoked (MD = -41.3%, 95% confidence interval [CI] -55.4% to -27.2%, n = 21, Figure 3) and ongoing (MD = -56.5%, 95%CI -75.3% to -37.6%, n = 14, Figure 4) activity in the EMTVN.

**Figure 3.**
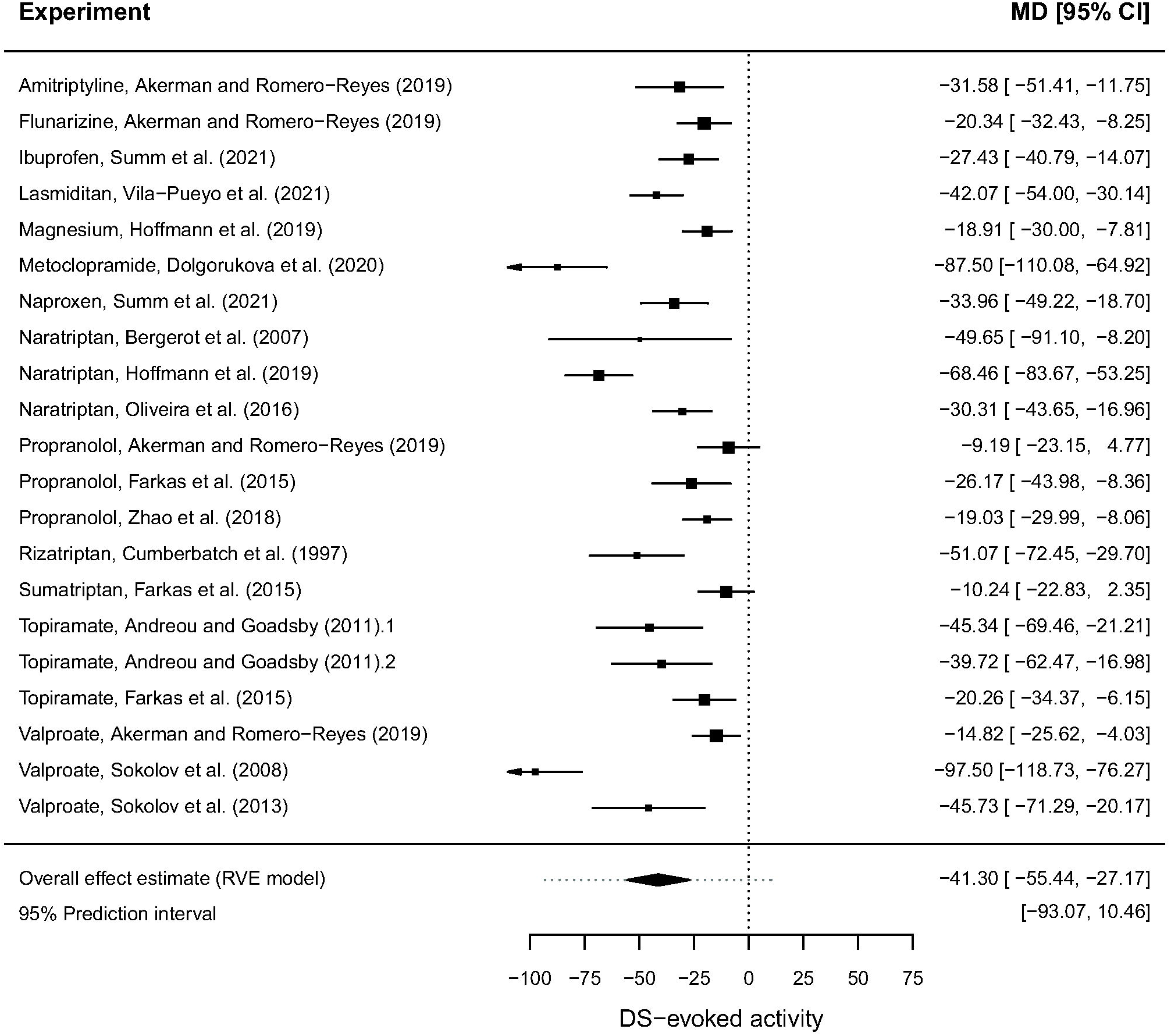
Forest plot of the 21 experiments reporting the drugs’ effects on the dural stimulation-evoked activity from the 13 studies (1-4 experiments per study, mean = 1.6) included in the analysis. The overall effect was calculated using robust variance estimation (RVE). MD, mean difference between treatment and control in %change of neuronal activity from baseline, CI, confidence interval.

**Figure 4.**
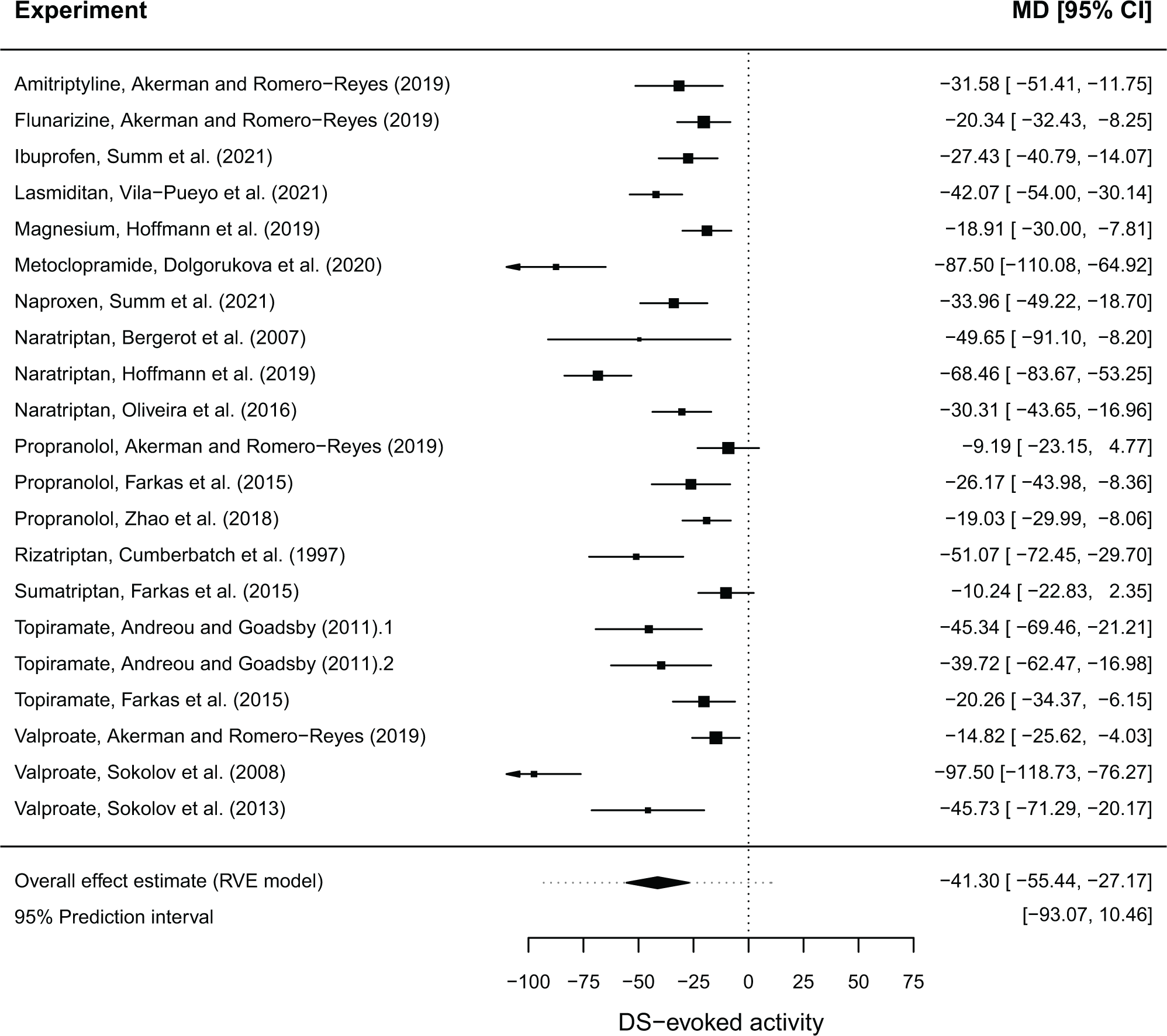
Forest plot of the 14 experiments reporting the drugs’ effects on the ongoing activity from the 8 studies (1-4 experiments per study, mean = 1.8) included in the analysis. The overall effect was calculated using robust variance estimation (RVE). MD, mean difference between treatment and control in %change of neuronal activity from baseline, CI, confidence interval.

In the ongoing activity data set, 64% of the variance was attributed to the overall heterogeneity. Between- and within-study heterogeneity accounted for 54% and 10% of the total heterogeneity, respectively. The remaining 36% was sampling variance. Since overall heterogeneity was only moderate, we did not perform subgroup and sensitivity analyses.

In the DS-evoked activity data set, the overall heterogeneity was 90% (between-study heterogeneity was 59%, and within-study heterogeneity was 31%). The remaining 10% was sampling variance. The extent to which methodological features explain the observed heterogeneity we assessed with meta-regression using the 6 factors predefined in the study protocol for grouping (the 7^th^ factor, sex of the animal, could not be analysed since all experiments were carried out in male rats). The DS-evoked activity inhibition was significantly more pronounces in experiments performed on room-air ventilated rats (MD = -78%, 95%CI -120% to -37%, n = 3) compared with O^2^-enriched air ventilated rats (MD = -29%, 95%CI -39% to -20%, n = 17; F_(1, 10)_ = 11.73, p = 0.006) (Supp. 3, Figure 3). Other factors did not account for a significant outcome variability (Supp. 3, Table 3).

The high heterogeneity in the data set can also be caused by extreme effect sizes, while the pooled effect estimate can be highly dependent on a single experiment. According to sensitivity analysis, 2/21 experiments from 2 studies (Dolgorukova et al., 2020; Sokolov et al., 2008) were influential outliers (Supp. 3, Figures 4-5). The results before and after the removal of the influential outliers are presented in Table 4.

**Table 4.**
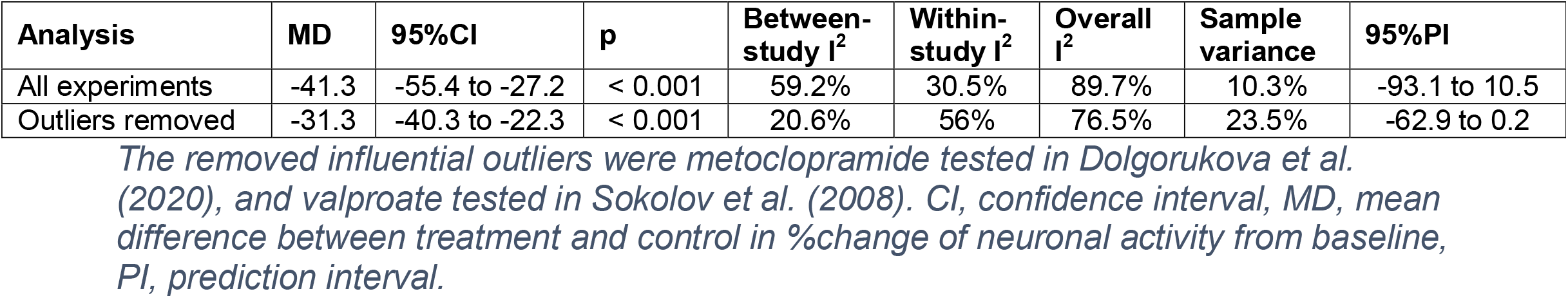
The results before and after the removal of the influential outliers.

Since after the exclusion of the two experiments overall heterogeneity was still high, we rerun subgroup analyses (Supp. 3, Table 5). The effect of gas mixture used for artificial ventilation was confirmed in this data set (F_(1, 8)_ = 14.82, p = 0.005). Besides, we revealed that the effect size estimate was significantly associated with the anaesthetic used for anaesthesia induction (F_(2, 8)_ = 4.83, p = 0.042), varying from smaller to larger in subgroups where were used urethane with or without alpha-chloralose (MD = -25%), sodium pentobarbital (MD = -30%), and inhalation anaesthetics (MD = -46%) (Supp. 3, Figures 6-7). Both factors, however, could not explain explicitly the overall heterogeneity (I^2^ = 77% and 75% for analyses where gas mixture and anaesthesia induction were used as moderators). To examine more generally the effect of approaches to EMTVN implementation in different research groups (the effect of laboratory), we additionally conducted a network analysis and revealed that there are four distinct clusters formed by study’s authors (Figure 5).

**Table 5.**
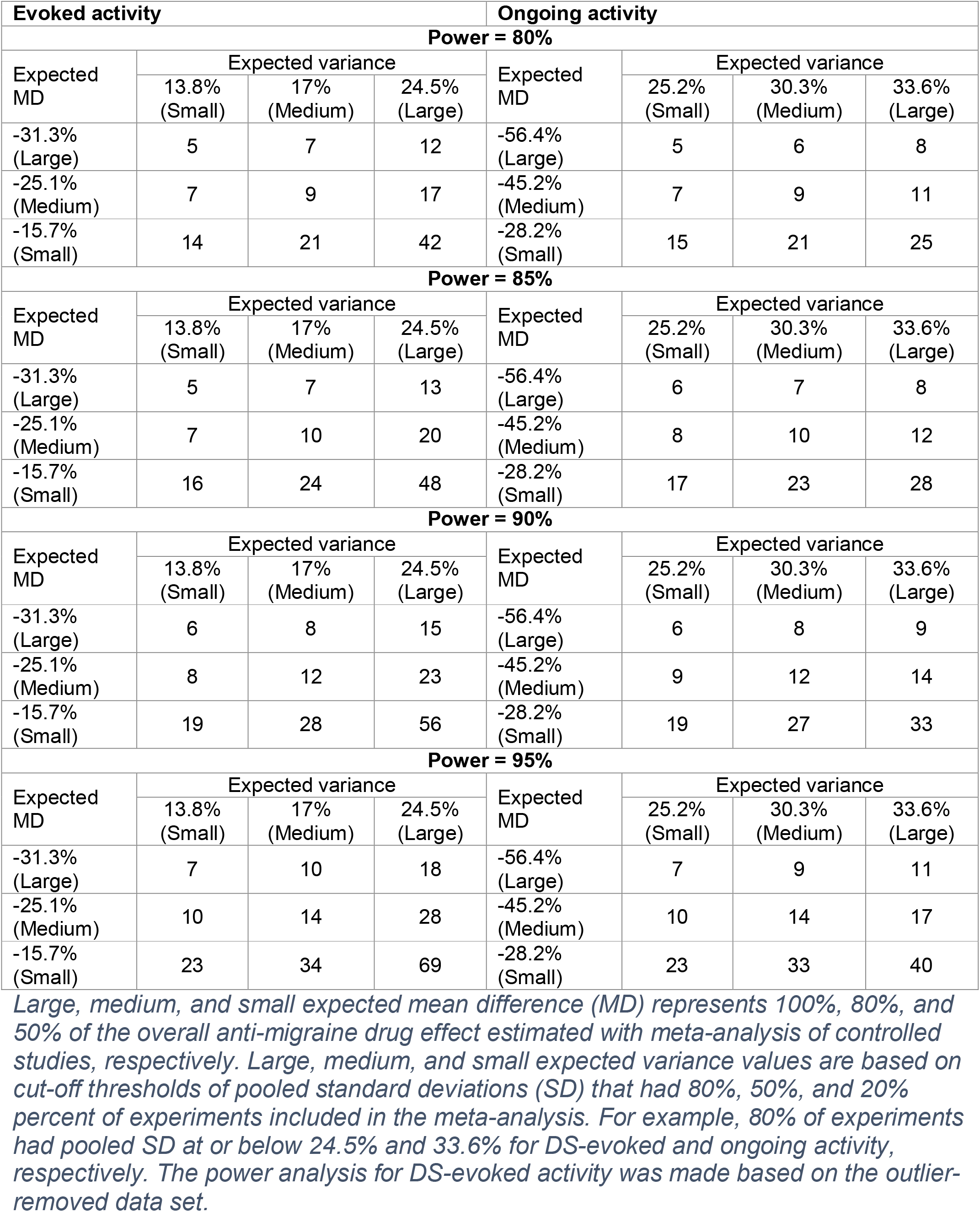
The number of animals per group required to detect the specified effects of anti-migraine drugs in the EMTVN with 80-95% power.

| Evoked activity |  |  |  | Ongoing activity |  |  |  |
| --- | --- | --- | --- | --- | --- | --- | --- |
| Power = 80% |  |  |  |  |  |  |  |
| Expected MD | Expected variance |  |  | Expected MD | Expected variance |  |  |
|  | 13.8% (Small) | 17% (Medium) | 24.5% (Large) |  | 25.2% (Small) | 30.3% (Medium) | 33.6% (Large) |
| -31.3% (Large) | 5 | 7 | 12 | -56.4% (Large) | 5 | 6 | 8 |
| -25.1% (Medium) | 7 | 9 | 17 | -45.2% (Medium) | 7 | 9 | 11 |
| -15.7% (Small) | 14 | 21 | 42 | -28.2% (Small) | 15 | 21 | 25 |
| Power = 85% |  |  |  |  |  |  |  |
| Expected MD | Expected variance |  |  | Expected MD | Expected variance |  |  |
|  | 13.8% (Small) | 17% (Medium) | 24.5% (Large) |  | 25.2% (Small) | 30.3% (Medium) | 33.6% (Large) |
| -31.3% (Large) | 5 | 7 | 13 | -56.4% (Large) | 6 | 7 | 8 |
| -25.1% (Medium) | 7 | 10 | 20 | -45.2% (Medium) | 8 | 10 | 12 |
| -15.7% (Small) | 16 | 24 | 48 | -28.2% (Small) | 17 | 23 | 28 |
| Power = 90% |  |  |  |  |  |  |  |
| Expected MD | Expected variance |  |  | Expected MD | Expected variance |  |  |
|  | 13.8% (Small) | 17% (Medium) | 24.5% (Large) |  | 25.2% (Small) | 30.3% (Medium) | 33.6% (Large) |
| -31.3% (Large) | 6 | 8 | 15 | -56.4% (Large) | 6 | 8 | 9 |
| -25.1% (Medium) | 8 | 12 | 23 | -45.2% (Medium) | 9 | 12 | 14 |
| -15.7% (Small) | 19 | 28 | 56 | -28.2% (Small) | 19 | 27 | 33 |
| Power = 95% |  |  |  |  |  |  |  |
| Expected MD | Expected variance |  |  | Expected MD | Expected variance |  |  |
|  | 13.8% (Small) | 17% (Medium) | 24.5% (Large) |  | 25.2% (Small) | 30.3% (Medium) | 33.6% (Large) |
| -31.3% (Large) | 7 | 10 | 18 | -56.4% (Large) | 7 | 9 | 11 |
| -25.1% (Medium) | 10 | 14 | 28 | -45.2% (Medium) | 10 | 14 | 17 |
| -15.7% (Small) | 23 | 34 | 69 | -28.2% (Small) | 23 | 33 | 40 |

*Large, medium, and small expected mean difference (MD) represents 100%, 80%, and 50% of the overall anti-migraine drug effect estimated with meta-analysis of controlled studies, respectively. Large, medium, and small expected variance values are based on cut-off thresholds of pooled standard deviations (SD) that had 80%, 50%, and 20% percent of experiments included in the meta-analysis. For example, 80% of experiments had pooled SD at or below 24.5% and 33.6% for DS-evoked and ongoing activity, respectively. The power analysis for DS-evoked activity was made based on the outlier-removed data set.*

**Figure 5.**
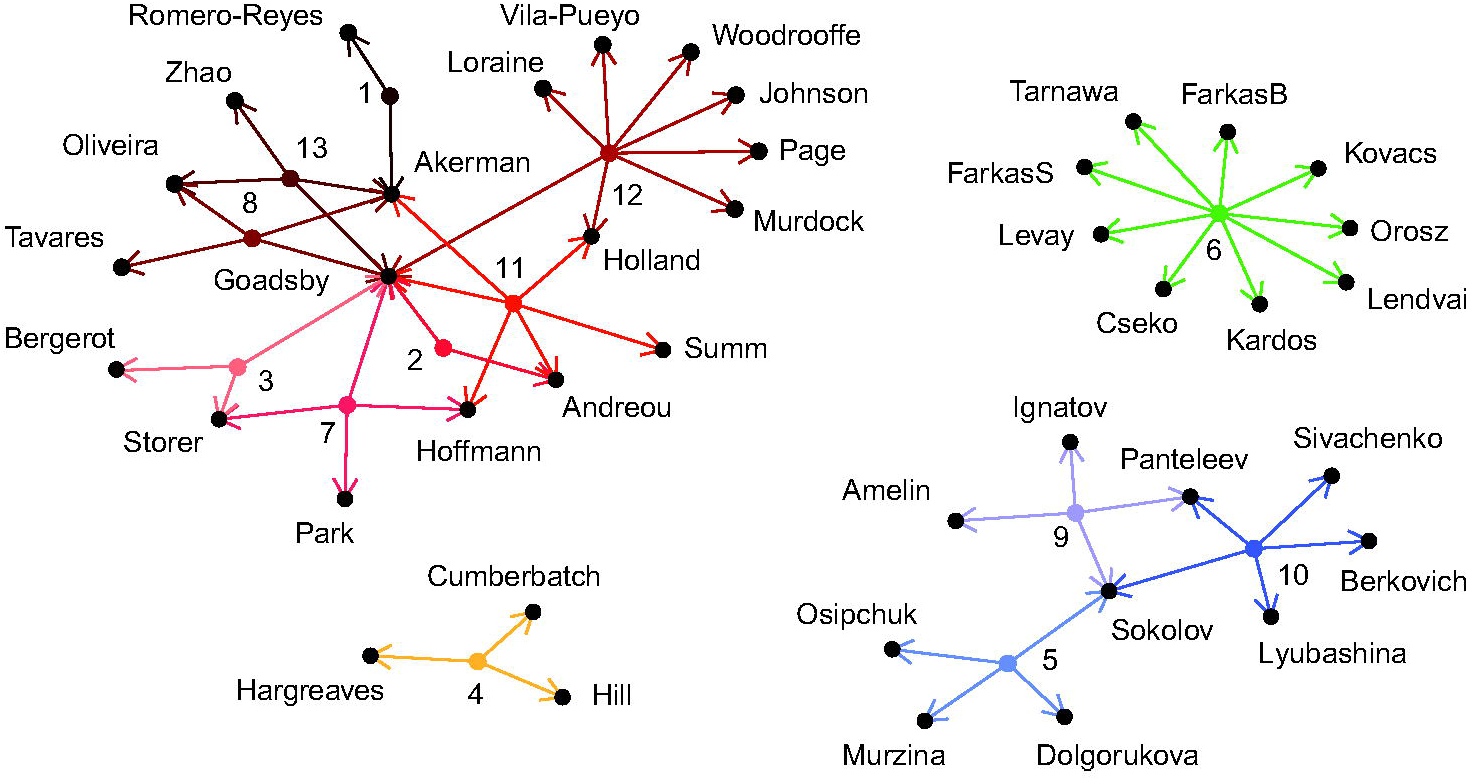
Networks analysis of study’s authors. The numbers represent study ID.

**Figure 6.**
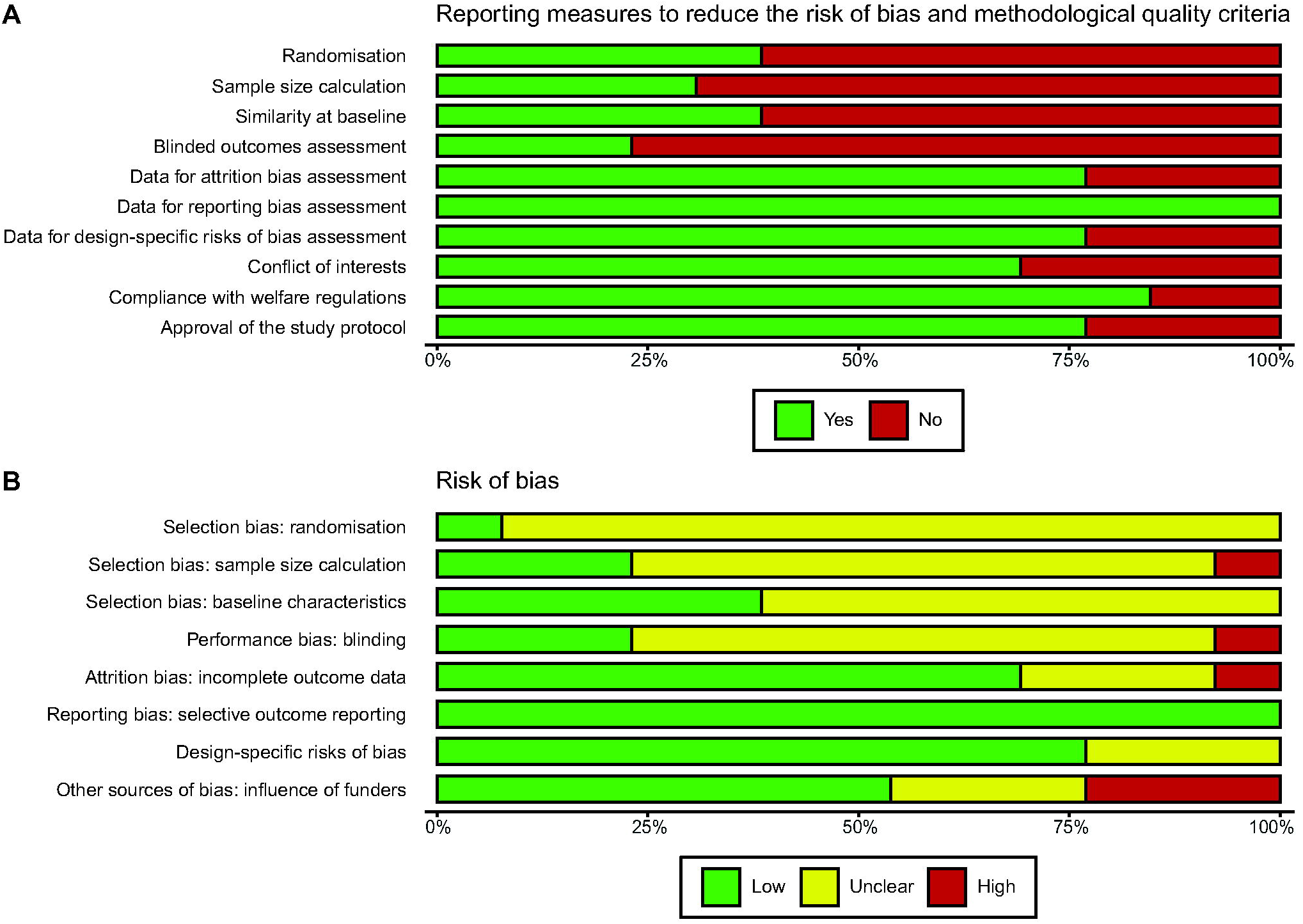
Summary bar plots showing the proportion of studies reporting measures to reduce risk of bias and methodological quality criteria (A) and the proportion of studies at low, high, and unclear risk of bias for each criteria (B) for the 13 included studies. Please, note that the two outcomes measured in these studies may have different risk assessments (i.e. attrition, design-specific risk of bias) but this was not the case, thus, all domains are presented at the study level. The “baseline characteristics” domain assesses reporting the absence of between-group baseline differences and/or accounting for baseline activity fluctuations in the analysis.

### 3.5. Risk of bias

The reporting of measures to reduce the risk of bias in the 13 included studies (Figure 6, A) was moderate: 38.5% (n = 5) reported randomisation to groups, 30.8% (n = 4) reported sample size calculation, 38.5% (n = 5) reported the absence of between-group difference at baseline and/or accounting for baseline fluctuations in the analysis (in 1 study baseline values were treated as covariant), 23.1% (n = 3) reported blinded assessment of outcomes and 1 study reported that the blinding was not possible due to noticeable treatment effect on blood pressure (assigned high risk of bias, Figure 6, B).

Overall, 5 studies (38.5%) did not report any of the above measures to reduce the risk of bias, and only one (7.7%) reported all four (Supp. 3, Figure 9).

The method of randomisation was rarely described: one of the 5 studies reporting randomisation used a Microsoft Excel random number generator, while the remaining studies did not report the method used. Across the 4 studies reporting sample size calculation, one used the resource equation method, and in two studies the sample size was calculated based on the estimated effect size. One publication reported that sample sizes were determined based on previous experience, which we do not consider a valid method for sample size calculation (assigned high risk of bias, Figure 6, B).

Much greater proportions of studies reported methodological quality criteria (Figure 6, A): 10 (76.9%) articles provided the data needed for attrition bias assessment (number of units tested and number of units for which the extracted outcomes are reported), one of them reported exclusion of a non-responder from the analysis, thus, we considered this study to be at high risk of attrition bias; all the included studies (100%) reported the data required for reporting bias assessment (all measured outcomes were specified in the methods section of the article).

The conflict of interest was stated in 9 (69.2%) publications (Figure 6, A). Based on these statements or, if absent, on author affiliations, 3 (23.1%) studies were considered at high risk of bias (the authors were affiliated with the pharmaceutical companies that synthesized the studied drugs, Figure 6, B).

In 10 (76.9%) studies, we did not reveal any design-specific risks of bias; in the remaining 3 publications, this risk was unclear (study design and/or the use of biological replicates were not clearly described) (Figure 6, B). The traffic light plot presenting the risk of bias assigned for each included study can be found in Supp. 3 (Figure 11).

Compliance with the international (e.g. EU Directive) or national animal welfare regulations was reported in 11 (84.6%) studies, and approval of the study protocol by an institutional animal care and use committee (or equivalent) in 10 (76.9%).

We did not reveal any associations between reporting measures to reduce the risk of bias and the drug effect on the DS-evoked activity. However, in the subset without influential outliers there was a significant difference between experiments reporting and not reporting randomisation (MD = -23% and -39%, respectively; F_(1, 9)_ = 7.15, p = 0.025) (Supp. 3, Tables 6-7). Of note, the outcome variability was lesser in the experiments, reporting randomisation (narrower confidence intervals, Supp. 3, Figure 13).

### 3.6. Publication Bias

Only the DS-evoked activity data set had a sufficient number of comparisons (≥ 20) to assess a potential publication bias. According to Egger’s regression test, there was a significant funnel plot asymmetry (p = 0.033) (Figure 7). We could not rerun this analysis without the influential outliers since this dataset include only 19 experiments (≥ 20 needed).

**Figure 7.**
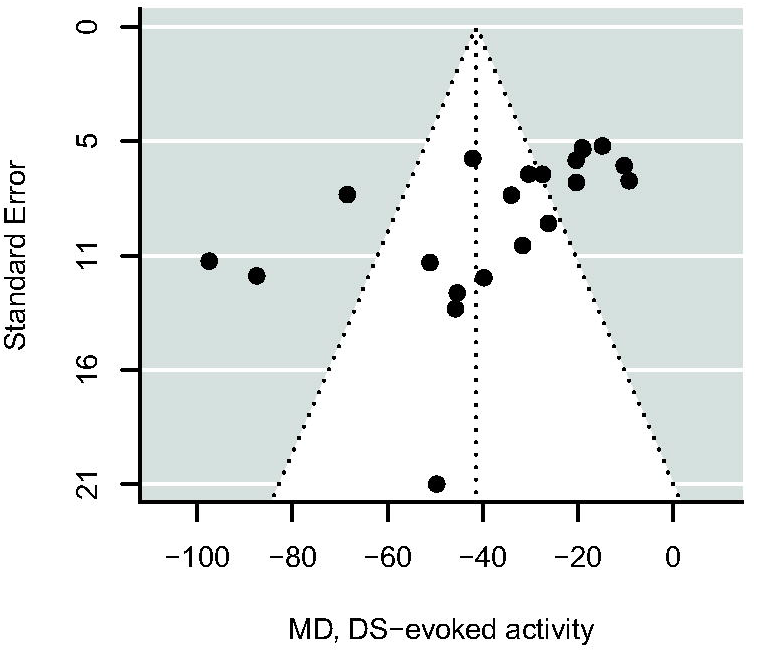
Funnel plot of the 21 experiments reporting the drugs’ effects on the dural stimulation-evoked activity from the 13 studies.

### 3.7. Sample size calculation

According to our calculations, to achieve 80-95% power, the detection of the effect of the same size as that of clinically effective anti-migraine drugs previously tested in controlled studies using the EMTVN may require 5 to 18 animals per group (Table 5). The exact number will depend on what outcome to measure, the expected outcome variance, and the desired study power.

For instance, in a lab where the DS-evoked activity showed a medium variation of around ±17% in previous or pilot experiments, detecting its 31% reduction compared to control at 80-85% power would require 7 animals per group (Table 5).

## 4. Discussion

Our systematic search identified 28 experiments, reported in 18 records, testing clinically effective anti-migraine drugs in the EMTVN. According to meta-analysis conducted for 21 extractable experiments (13 studies), these treatments significantly decrease trigeminovascular nociceptive traffic compared with a control intervention. The estimated effects were used as reference effect sizes to calculate the sample sizes allowing to achieve sufficient power in future experiments. In the relevant sections below, we discuss the translational potential of the EMTVN and what bias should be addressed for its improvement.

### 4.1. External validity of the EMTVN

External validation estimates the extent to which the results can be translated to a population with the modelled disease (O’Connor and Sargeant, 2014). It is usually made using three main criteria: construct, face, and predictive validity.

The similarity of the TVS anatomy and function in humans and animals, as well as the strong theoretical rationale behind the EMTVN, yield convincing construct validity. At the same time, since it is performed under general anaesthesia, it is completely devoid of face validity, which is a recapitulation of the disease phenotype. The third and most important criterion for external validation of an animal model in drug development, predictive validity, has been demonstrated previously and confirmed in the present study.

According to research data, substances exacerbating migraine pain as well as migraine triggers significantly enhance neuronal excitability in the EMTVN (Akerman et al., 2019; Burstein et al., 1998; Messlinger et al., 2010; Noseda et al., 2010). At the same time, clinically effective non-pharmacological manipulations (Andreou et al., 2016; Lyubashina et al., 2012) and medications (the present results) suppress trigeminovascular nociceptive traffic. This is primarily manifested by the inhibition of neuronal responses to stimulation of the dura mater and/or the ongoing neuronal activity.

Regarding the first outcome, the stimulation-evoked activity, it should be noted that meningeal sensory innervation is represented mainly by Aδ- (2-30 m/s) and C-type (< 2 m/s) fibres, the responses of which can be differentiated in the EMTVN. However, in most of the included studies, neurons with only Aδ-fibre, mixed, or undisclosed inputs were recorded. Also, we combined the Aδ- and C-fibre response data that were separately reported in the remaining studies and, therefore, could not assess how much each of them contributed to the predictive value of the evoked activity inhibition. The second outcome in the EMTVN is the ongoing firing rate. It consists of the intrinsic discharges of trigeminovascular neurons as well as those that result from the spatiotemporal summation of impulses arriving from the periphery, e.g. from skin and muscles damaged during surgical preparation (Nöbel et al., 2016; Roch et al., 2007) and is under control of descending modulation. Given that the ongoing activity despite its origin belongs to specific trigeminovascular neuronal population, its reduction, as that of the evoked activity, is considered as a marker of potential antimigraine action. It has also been hypothesised that the intrinsic firing rate of trigeminovascular neurons may correlate with their susceptibility to migraine triggers and, thus, its dampening could predict a drug’s ability to reduce the frequency of migraine attacks (Akerman and Romero-Reyes, 2019). As more data on migraine preventives becomes available, it will be interesting to evaluate this hypothesis further. Another question left for future research is whether the antimigraine drugs’ effects on the evoked and ongoing activity are comparable. Since we have performed separate analyses for each outcome, we could not examine it here.

While the overall significant inhibitory effect of clinically effective anti-migraine drugs on the ongoing and evoked neuronal activity indicates predictive validity of the EMTVN, robustness of the results may be of some concern. According to studies’ reports, the same drug may demonstrate considerably varying effect from one experiment to another. For instance, 30 min after administration, naratriptan at 10 mg/kg inhibited the evoked neuronal activity by 30% of baseline in Oliveira et al. (2016) and about twice as much in Hoffmann et al. (2019), despite the fact that it was used at a lower dose of 5 mg/kg. This may be a result of great between-animal variability of this outcome, which would indicate that only the direction of the effect matters in the EMTVN. Besides, this may be due to other factors, such as different experimental conditions or statistical bias.

### 4.2. Possible sources of variability in the outcomes measured in the EMTVN

The inhibitory effect on the ongoing neuronal firing turned out to be more variable (wider confidence intervals) but also more consistent between the experiments and studies based on moderate overall heterogeneity. We, therefore, were not able to determine whether alterations in methodology affected the magnitude of the ongoing activity decrease. In contrast, the heterogeneity in drug effects on the neuronal activity evoked by dural stimulation was high even after the removal of two influential outliers from this data set. Consequently, we performed subgroup analyses with and without influential outliers.

According to our results, the magnitude of the evoked activity inhibition may vary depending on the gas mixture (O_2_-enriched or room air) used for the artificial ventilation. This may be due to the direct confounding effect of oxygen (e.g. its vasoconstrictive action, tissue oxygenation level) or due to other factors dividing the experiments into the same subgroups as the gas mixture, which were not evaluated in this study (e.g. ventilation parameters, laboratory equipment, geographical region, etc.). Interestingly, the smaller estimate of the effect was for the experiments performed in rats ventilated with O_2_-enriched air. In fact, oxygen can be involved in migraine pathogenesis in different ways that have been reviewed elsewhere (Bennett et al., 2015; Ciarambino et al., 2021; Taylor, 2011), although in the EMTVN, 100% oxygen had no direct effect on the neuronal responses to dural stimulation (Akerman et al., 2009). At the same time, lowering O_2_ in the ventilation gas was accompanied by transient activation of the trigeminovascular neurons, while the prolonged hypoxia led to its progressive decrease or even termination (Waldmann and Messlinger, 2021). This cannot, however, explain the greater reduction of neuronal excitability under ventilation with room air, which rather provides normoxic conditions. Thus, it would be interesting to further explore the possible influence of the gas mixture in the EMTVN experimentally.

The other interesting finding is that anaesthetics can also affect the outcome in the EMTVN. We have detected the effect of anaesthesia induction but not maintenance, although the latter may be due to the fact that different regimens were used (it is usually administered as required), which could cause varying effect. There were three groups of anaesthetics used for anaesthesia induction: urethane with or without alpha-chloralose, sodium pentobarbital, and inhalation anaesthetics, with the first group giving the smallest estimate of the effect and the last group giving the greater. This may be due to varying degree of analgesia produced by these drugs or due to their different receptor binding affinities and influence on excitatory/inhibitory transmission (Evgenov et al., 2020; Flecknell, 2016a, b; Vuyk et al., 2020).

It should be noted that after including these two factors as moderators, the residual heterogeneity remained high, suggesting that there are other moderators not considered in the model. Our additional network analysis has shown that the authors of the included publications represent four distinct research groups. Thus, the influence of both factors (gas mixture and anaesthesia induction) can also be a consequence of different approaches to EMTVN implementation in these research groups (the effect of the laboratory). The network analysis we have used here can serve as a tool for identifying distinct research groups in subgroup analyses in future works.

### 4.3. Internal validity of studies

Internal validity refers to how the study results represent the source population (Huang et al., 2020; O’Connor and Sargeant, 2014). It can be assessed by reviewing what steps have been taken to control possible systematic errors.

Overall, 5 of 13 studies (38.5%) reported no measures to reduce the risk of bias and only 1 (7.7%) reported all four (randomisation, sample size calculation, similarity at baseline, and blinding). Surprisingly few studies (n = 5, 38.5%) reported randomisation and the absence of between-group difference at baseline and/or accounting for baseline fluctuations in the analysis, which we believe is fundamental for between-group comparisons. The reporting of sample size calculation and blinding was also exceptional. Unfortunately, due to poor reporting on these protective measures, we were unable to assess the risk of bias reliably, which compromises our findings.

Several systematic reviews have shown that not reporting measures to reduce the risk of bias in biomedical research is associated with larger effect sizes (Hirst et al., 2013; Macleod et al., 2008; Vesterinen et al., 2010), but the opposite effect has also been demonstrated (Egan et al., 2016; Soliman et al., 2021). This discrepancy suggests that the prime source of systematic error is different for each animal model. For instance, in the EMTVN the data are gathered using objective assessment, which is expected to be less prone to biases. On the other hand, animals are usually tested one per day, and the experimental set can take months. Without randomisation, differences in baseline characteristics between treatment groups are very likely, indicating that the selection bias may have the strongest impact on the outcome. Indeed, for the experiments included in this review, the reporting of randomisation was associated with smaller effect sizes. Of note, this effect became apparent only after the removal of influential outliers from the data set. Although we have not identified other sources of bias, this should not be interpreted as the absence of such.

Much greater proportions of studies (69 – 100%) reported methodological quality criteria, such as the number of units tested and the number of units for which the results are reported, the measured outcomes, clear description of the design used, compliance with animal welfare regulations, approval of the study protocol, and conflict of interest statement.

### 4.4. Publication bias

Only the evoked activity data set had a sufficient number of comparisons (≥ 20) to assess the potential publication bias. The funnel plot inspection, supported by Egger’s regression test, indicates a clear trend towards increasing effect size with decreasing precision. This may be simply due to the presence of missing unpublished experiments with similar standard errors but null or negative results (publication bias). Another possible reason is that experiments with lower precision are likely to be of poor quality and therefore at higher risk of bias (Button et al., 2013; Egger et al., 1997; Page et al., 2021). In light of the fact that studies not reporting randomisation yielded effects of greater magnitude, this is a more likely explanation. Finally, funnel plots do not account for confounding variables, and the high heterogeneity as well as biological variance may also cause the funnel plot asymmetry (Egger et al., 1997; Page et al., 2021).

Either way, the overall effects of the anti-migraine drugs are likely overestimated in this review, supplementing the previous report, according to which animal studies on neurological disorders are prone to excess significance (Tsilidis et al., 2013).

### 4.5. Sample size

Sample size calculation is a pivotal aspect of planning a preclinical study, but it is complicated by defining of an effect size to be detected, which should be biologically relevant. As we have outlined in the introduction, here we have hypothesised that the summary effect of previously tested clinically effective drugs may be considered biologically relevant and support sample size calculation. Our meta-analysis indicated that in the EMTVN, such drugs exhibit the mean difference of 31% (excluding outlying experiments) and 56% compared to control intervention for the DS-evoked and ongoing activity, respectively. Consequently, the EMTVN requires from 5 to 18 animals per group depending on the expected outcome variance and the desired study power.

The summary effect derived with meta-analysis, however, is likely overestimated. Consequently, we provide guidance on sample sizes allowing sufficiently power a study for detection 80% and 50% of the reference effect.

The selection of the effect size and power to achieve should be based on the study aims, previous data, and the design of the experiments. The latter is particularly important, since there are at least two potential confounders related to EMTVN implementation (gas mixture for artificial ventilation and anesthetics). Besides, here we show that the effect size and variability in the data may also depend on randomisation. Therefore, we suggest keeping these factors in mind while planning study design and using our calculations to get an idea of the effect sizes that one could reasonably expect to demonstrate with given sample size, data variability, and power. More guidance on sample size calculation can be found in NC3R website (Experimental design assistant, https://eda.nc3rs.org.uk/experimental-design-group#samplesize).

Of note, when the outcome variance is expected to be large, we recommend preliminary design optimization, as the reduction of variability, especially of DS-evoked activity, may lead to essential decrease of sample sizes. This may include randomisation and other measures to reduce risk of bias; local anaesthesia during surgical preparation (Laborc et al., 2020) and provision of a habituation period thereafter; equipment setup allowing to reduce background noise and interferences (Faraday cage, spike sorting and filtration, etc.) and improving the quality of neuronal activity registration.

### 4.6. Limitations

The main limitation of a meta-analysis is that it is based on the information that has been reported in publications. Also, the list of included experiments in this work is not exhaustive, since there were several studies with missing information that prevented data extraction for a quarter of experiments (7 of 28 experiments in 4 of the 17 included studies) and several studies, from which we were only able to extract a subset of the experiments. Besides, we have detected a trend towards increasing effect size with decreasing precision and poor reporting on measures taken to reduce the risk of systematic error. The drugs’ effects estimated with meta-analyses can, thus be not accurate. However, in addressing the principal question of this work, we took into account the possible effect size overestimation and provided power analysis allowing the detection of 80% and 50% of the overall effect sizes. It is worth noting that all the included experiments were performed in male rats, thus our guidance on sample sizes should not be generalized to female rats or other species.

The analyses include unpublished data provided by the authors, while some authors of this work have also authored several included studies. However, we explicitly stated the unpublished additions and made available both the raw data and program codes allowing the reproduction of the results.

Finally, the findings that different approaches to EMTVN implementation can impact the results obtained in this model should be interpreted with caution since they were revealed in explorative analyses and require experimental confirmation. Besides, the high amount of heterogeneity remains unexplained and should be addressed in further updates.

## Supporting information

Supplement file 1: Search terms

Supplement file 2: The characteristics of the included studies and experiments

Supplement file 3: Analysis

## 5. Data availability

The raw and processed data, R Markdown scripts producing the results as well as study protocol and all supplemental materials can be found at the project page: https://osf.io/vzjys/?view_only=9c7a38db19304c94b38e1fc805c470aa

## 6. Acknowledgments

The authors wish to express their gratitude to the researchers without whom this work would not have been possible but who preferred not to be named. The authors acknowledge the fruitful efforts of research groups elaborating open-source materials and frameworks, which substantially facilitate systematic reviews of animal studies (The Collaborative Approach to Meta-Analysis and Review of Animal Experimental Studies: CAMARADES). Finally, the authors are grateful to the developers of SYRF - the platform supporting the data extraction, and the metafor package for R, who provide valuable guidance on these tools’ implementation in meta-analyses.

## 8. Author Contributions

Conceptualisation: Antonina Dolgorukova, Alexey Y. Sokolov, Elena Verbitskaya. Screening of abstracts and full text: Antonina Dolgorukova, Julia Isaeva, Ekaterina Protsenko, Victoria Gagloeva. Data extraction: Antonina Dolgorukova, Ekaterina Protsenko, Victoria Gagloeva. Data reconciliation: Antonina Dolgorukova, Victoria Gagloeva, Alexey Y. Sokolov. Analysis and interpretation of the data: Antonina Dolgorukova, Alexey Y. Sokolov, Elena Verbitskaya; Writing – original draft: Antonina Dolgorukova; Writing – review & editing: Ekaterina Protsenko, Julia Isaeva, Victoria Gagloeva, Elena Verbitskaya, Alexey Y. Sokolov. Final approval of the manuscript: all authors.

