## Supplement file 1: Search terms for "Meta-analysis of the effects of clinically-effective therapeutics in the preclinical migraine model as a tool for design optimisation"

Antonina Dolgorukova, Ekaterina Protsenko, Julia Isaeva, Victoria Gagloeva, Elena Verbitskaya, Alexey Y. Sokolov.

### 1. Pubmed

((trigeminovascular[All Fields] OR craniovascular[All Fields] OR durovascular[All Fields]) AND ("nociception"[MeSH Terms] OR "nociception"[All Fields] OR activation[All Fields] OR "activity"[All Fields] OR response [All Fields] OR responses[All Fields])) OR (stimulation[All Fields] AND ((middle[All Fields] AND meningeal[All Fields] AND ("arteries"[MeSH Terms] OR "arteries"[All Fields] OR "artery"[All Fields])) OR dura[All Fields] OR dural[All Fields] OR (sagittal[All Fields] AND sinus[All Fields]))) OR ((electrophysiological[All Fields] OR extracellular[All Fields] OR single-unit[All Fields] OR multi-unit[All Fields]) AND ( model[All Fields] OR models[All Fields] OR "techniques"[All Fields] OR technique[All Fields] OR "assay"[All Fields] OR recording[All Fields] OR recordings[All Fields] OR recorded[All Fields]) AND ((trigeminal[All Fields] OR trigeminocervical[All Fields] OR trigeminothalamic[All Fields] OR trigeminothalamocortical[All Fields] OR trigeminovascular[All Fields]) OR meningeal[All Fields]) AND ("neurons"[MeSH Terms] OR "neurons"[All Fields] OR "neuron"[All Fields] OR neurones[All Fields] OR neuronal[All Fields] OR "nociceptors"[MeSH Terms] OR "nociceptors"[All Fields] OR "nociceptor"[All Fields] OR afferent[All Fields] OR afferents[All Fields])) OR ((dura[All Fields] OR dural[All Fields] OR meningeal[All Fields]) AND ("nociceptors"[MeSH Terms] OR "nociceptors"[All Fields] OR "nociceptor"[All Fields] OR afferent[All Fields] OR afferents[All Fields] OR trigeminal[All Fields] OR trigeminocervical[All Fields] OR trigeminovascular[All Fields] OR (second[All Fields] AND order[All Fields]) OR second-order[All Fields] OR trigeminothalamic [All Fields] OR (third[All Fields] AND order[All Fields]) OR third-order[All Fields] OR thalamocortical[All Fields] OR trigeminothalamocortical[All Fields]) AND ("neurons"[MeSH Terms] OR "neurons"[All Fields] OR "neuron"[All Fields] OR neurones[All Fields]))

Filter: publication date from 1986

26/09/21 2,771 results

14/12/21 2,793 results

### 2. Scopus

TITLE-ABS-KEY ( stimulat* AND "middle meningeal arter*" OR dura OR dural OR "sagittal sinus" ) ) OR ( TITLE-ABS-KEY ( ( trigeminal OR trigeminocervical OR trigeminothalamic OR trigeminovascular OR meningeal OR craniovascular ) AND ( nociception OR activation OR activity ) AND ( migraine OR headache ) AND ( model OR technique OR assay OR record* ) ) ) OR ( TITLE-ABS-KEY ( ( ( electrophysiological OR extracellular OR single-unit OR multi-unit ) AND ( model OR technique OR assay OR record* ) ) OR ( dura OR dural OR "middle meningeal arter*" OR "sagittal sinus" ) AND ( trigeminal OR trigeminocervical OR ( "second order" AND trigemin* ) OR trigeminothalamic OR trigeminothalamocortical OR ( "third order" AND trigemin* ) OR trigeminovascular OR meningeal OR craniovascular ) AND ( neuron OR neuronal OR nociceptor OR afferent ) ) ) AND PUBYEAR > 1985 / AND ORIG-LOAD-DATE AFT 20210926

26/09/21 3,219 results

14/12/21 22 new results

Study protocol is available at PROSPERO website (CRD42021276448; <https://www.crd.york.ac.uk/prospero/display_record.php?ID=CRD42021276448>)
