## Supplement file 2: The characteristics of the included studies and experiments for "Meta-analysis of the effects of clinically-effective therapeutics in the preclinical migraine model as a tool for design optimisation"

Antonina Dolgorukova, Ekaterina Protsenko, Julia Isaeva, Victoria Gagloeva, Elena Verbitskaya, Alexey Y. Sokolov.

The analysis included 13 studies assessing the effects of 13 anti-migraine drugs on the ongoing (8 studies, 14 experiments) and dural stimulation-evoked (DS-evoked, 13 studies, 21 experiments) activity of 290 neurons/neuronal clusters using the electrophysiological model of trigeminovascular nociception (EMTVN).

### 1. Studies

Table 1: Study locations

| Study.ID | Reference | Country | City | Organisation |
| --- | --- | --- | --- | --- |
| 1 | Akerman and Romero-Reyes (2019) | USA | New York | Department of Oral and Maxillofacial Pathology, Radiology and Medicine, New York University College of Dentistry |
| 2 | Andreou and Goadsby (2011) | USA | San Francisco | Department of Neurology, University of California |
| 3 | Bergerot et al. (2007) | UK | London | Institute of Neurology, and National Hospital for Neurology and Neurosurgery |
| 4 | Cumberbatch et al. (1997) | UK | Essex | Merck Sharp&Dohme Research Laboratories, Neuroscience Research Center |
| 5 | Dolgorukova et al. (2020) | Russia | Saint Petersburg | Valdman Institute of Pharmacology, Pavlov First Saint Petersburg State Medical University |
| 6 | Farkas et al. (2015) | Hungary | Budapest | Pharmacology and Drug Safety Research, Gedeon Richter Plc |
| 7 | Hoffmann et al. (2019) | USA | San Francisco | Department of Neurology, University of California |
| 8 | Oliveira et al. (2016) | UK | London | Basic and Clinical Neuroscience, Institute of Psychology, Psychiatry and Neuroscience, King’s College London |
| 9 | Sokolov et al. (2008) | Russia | Saint Petersburg | Valdman Institute of Pharmacology, Pavlov First Saint Petersburg State Medical University |
| 10 | Sokolov et al. (2013) | Russia | Saint Petersburg | Laboratory of Cortico-Visceral Physiology, Pavlov Institute of Physiology of the Russian Academy of Sciences |
| 11 | Summ et al. (2021) | USA | San Francisco | Department of Neurology, University of California |
| 12 | Vila-Pueyo et al. (2021) | UK | London | Wolfson Centre for Age-Related Diseases, Institute of Psychiatry, Psychology & Neuroscience, King’s College London |
| 13 | Zhao et al. (2018) | USA | San Francisco | Department of Neurology, University of California |

11 studies used parallel design and 2 used unclear design.

Inclusion and/or exclusion criteria (other than sensitivity to dural stimulation) were reported in 11/13 studies.

Table 2: Inclusion and exclusion criteria

| Study.ID | Reference | Inclusion.and.exclusion.criteria |
| --- | --- | --- |
| 1 | Akerman and Romero-Reyes (2019) | Neuronal clusters; input from the facial skin innervated by the first (ophthalmic) branch of the trigeminal nerve. |
| 2 | Andreou and Goadsby (2011) | A probability of neuronal firing of greater than 30%. |
| 3 | Bergerot et al. (2007) | NR |
| 4 | Cumberbatch et al. (1997) | The experiment terminated if the systolic pressure remained below 90 mmHg |
| 5 | Dolgorukova et al. (2020) | Input from the facial skin; stable and reproducible over time firing rate. |
| 6 | Farkas et al. (2015) | Neuronal clusters; WDR input from the facial skin innervated by the first (ophthalmic) branch of the trigeminal nerve; a mean ongoing activity between 5 and 50 Hz; animals exhibiting lower than 60 mmHg mean arterial blood pressure or higher than 50 mmHg fluctuation for a period longer than 5 minutes were excluded. |
| 7 | Hoffmann et al. (2019) | WDR input from the facial skin innervated by the first (ophthalmic) branch of the trigeminal nerve. |
| 8 | Oliveira et al. (2016) | Input from the facial skin innervated by the first (ophthalmic) branch of the trigeminal nerve. |
| 9 | Sokolov et al. (2008) | Input from the facial skin in the fronto-orbital-temporal region. |
| 10 | Sokolov et al. (2013) | Input from the facial skin. |
| 11 | Summ et al. (2021) | WDR input from the facial skin innervated by the first (ophthalmic) branch of the trigeminal nerve; a mean firing rate of 30% above baseline within a 7 to 10 ms period of the main firing episode (A delta fibres). |
| 12 | Vila-Pueyo et al. (2021) | Input from the facial skin innervated by the first (ophthalmic) branch of the trigeminal nerve. |
| 13 | Zhao et al. (2018) | NR |

*NR, not reported, WDR, wide dynamic range units (responsive to both noxious and non-noxious mechanical stimulation of cutaneous receptive fields).*

### 2. Animals

The included experiments (n = 21) were performed in male rats.

Table 3: Strains of the rats used

| Strain | n |
| --- | --- |
| Sprague-Dawley | 15 |
| Wistar | 6 |

The rats’ weight was within the range 220 - 440 g, and the age was either adult (n = 2 studies) or not reported (11 studies).

We could not calculate the number of animals used because:

- the number of animals per group was not reported in 4 experiments;
- it was not equal to the number of neurons recorded (n = 290 neurons/neuronal clusters), since 3 studies reported biological replicates (i.e when more than one neuron/cluster was recorded per animal). None of them reported whether the units were recorded simultaneously or sequentially.

### 3. Model characteristics

The anaesthetics were grouped as follows:

- urethane and alpha-chloralose or urethane only: Urethane with or without alpha-chloralose;
- isoflurane, halothane: Inhalation anesthetics.

Table 4: Anaesthetics used to initiate anaesthesia in 21 experiments with DS-evoked activity measurement and in 14 experiments with ongoing activity measurement

| Anaesthesia.induction | DS-evoked activity | Ongoing activity |
| --- | --- | --- |
| Inhalation anesthetics | 2 | 1 |
| Sodium pentobarbital | 13 | 7 |
| Urethane with or without alpha-chloralose | 6 | 6 |

*DS-evoked, dural stimulation-evoked.*

Table 5: Anaesthetics used for anaesthesia maintenence in 21 experiments with DS-evoked activity measurement and in 14 experiments with ongoing activity measurement

| Anaesthesia.maintenance | DS-evoked activity | Ongoing activity |
| --- | --- | --- |
| Propofol | 10 | 6 |
| Sodium pentobarbital | 5 | 2 |
| Urethane with or without alpha-chloralose | 6 | 6 |

*DS-evoked, dural stimulation-evoked.*

Artificial ventilation was used in 13/13 studies.

Table 6: Gas mixture used for artificial ventilation in 21 experiments with DS-evoked activity measurement and in 14 experiments with ongoing activity measurement

| Gas.mixture | DS-evoked activity | Ongoing activity |
| --- | --- | --- |
| NR | 1 | 0 |
| O2-enriched air | 17 | 11 |
| Room air | 3 | 3 |

*DS-evoked, dural stimulation-evoked, NR, not reported.*

The use of a myorelaxant was reported in 8/13 studies.

The use of other drugs during surgical preparation was reported in 1 study (atropine, 50 ug/kg, s.c.; acetazolamide, 10 mg/kg, i.p.; lignocaine, 10% w/w, was used routinely on skin incisions, in the region of femoral artery cannulation, and in the ear canal before mounting in the stereotaxic apparatus)

Studies used electrical stimulation of the meninges to activate trigeminovascular neurons.

Table 7: Stimulation parameters

| Study.ID | Reference | Stimulation.parameters | n |
| --- | --- | --- | --- |
| 1 | Akerman and Romero-Reyes (2019) | 4-15 V, 0.1-0.2 ms, 0.25 Hz (Grass S88 stimulator) | 4 |
| 2 | Andreou and Goadsby (2011) | 50-150 uA, 0.1-0.2 ms, 0.5 Hz (Grass S88 stimulator) | 2 |
| 3 | Bergerot et al. (2007) | 15 V, 1 ms, 0.5 Hz (NE 200X) | 1 |
| 4 | Cumberbatch et al. (1997) | 2000-6000 uA, 0.07-0.5 ms, 1 Hz, every 200 s | 1 |
| 5 | Dolgorukova et al. (2020) | 15-40 V, 0.2-0.6 ms, 0.33 Hz (Master-8) | 1 |
| 6 | Farkas et al. (2015) | 1.5-5 V, 0.8 ms, 0.033 Hz (Biostim) | 3 |
| 7 | Hoffmann et al. (2019) | 8-22 V, 0.1-0.2 ms, 0.5 Hz (Grass S88 stimulator) | 2 |
| 8 | Oliveira et al. (2016) | 8-15 V, 0.15-0.25 ms, 0.4-0.5 Hz (Grass S88 stimulator) | 1 |
| 9 | Sokolov et al. (2008) | 500-1000 uA, 0.25 ms, 0.3 Hz (Universal Electrostimulator type UES-2) | 1 |
| 10 | Sokolov et al. (2013) | 15-30 V, 300-600 uA, 0.3-0.5 ms | 1 |
| 11 | Summ et al. (2021) | 5-16 V, 0.1-0.2 ms, 0.5 Hz | 2 |
| 12 | Vila-Pueyo et al. (2021) | 8-15 V, 0.3-0.5 ms, 0.5 Hz | 1 |
| 13 | Zhao et al. (2018) | 10-20 V, 0.1-0.3 ms, 0.5 Hz (Grass S88 stimulator) | 1 |

*n, number of experiments.*

Table 8: Stimulation intensity

| Study.ID | Reference | Stimulation.intensity | n |
| --- | --- | --- | --- |
| 1 | Akerman and Romero-Reyes (2019) | Threshold | 4 |
| 2 | Andreou and Goadsby (2011) | Threshold | 2 |
| 3 | Bergerot et al. (2007) | NR | 1 |
| 4 | Cumberbatch et al. (1997) | Surpathreshold | 1 |
| 5 | Dolgorukova et al. (2020) | Surpathreshold | 1 |
| 6 | Farkas et al. (2015) | Surpathreshold | 3 |
| 7 | Hoffmann et al. (2019) | NR | 2 |
| 8 | Oliveira et al. (2016) | Surpathreshold | 1 |
| 9 | Sokolov et al. (2008) | Surpathreshold | 1 |
| 10 | Sokolov et al. (2013) | Surpathreshold | 1 |
| 11 | Summ et al. (2021) | Threshold | 2 |
| 12 | Vila-Pueyo et al. (2021) | Threshold | 1 |
| 13 | Zhao et al. (2018) | Surpathreshold | 1 |

*NR, not reported, n, number of experiments.*

Habituation period (period of time during which animals rested after surgical preparation) was reported in 3/13 studies and was 30 to 60 min.

Overall, 290 neurons or neuronal clusters were recorded in the trigeminocervical complex (TCC, 12 studies, 19 experiments) or ventroposteromedial thalamic nucleus (2 studies, 2 experiments).

3 studies used multi-unit recording (i.e when more than one neuron/cluster was recorded per animal), and none of them reported whether the units were recorded simultaneously or sequentially. 10 studies used single-unit recording (1 unit per animal). None of the included studies reported spike sorting or filtration after the recordings.

### 4. Recorded neurons

Table 9: Coordinates of TCC neurons: range

| Study.ID | Reference | Recording site | AP min | AP max | ML min | ML max | DV min | DV max |
| --- | --- | --- | --- | --- | --- | --- | --- | --- |
| 1 | Akerman and Romero-Reyes (2019) | TCC | NR | NR | NR | NR | 0.10 | 1.10 |
| 4 | Cumberbatch et al. (1997) | TCC | 1 | 2 | NR | NR | 0.40 | 1.21 |
| 5 | Dolgorukova et al. (2020) | TCC | 0.94 | 4.56 | 0.74 | 1.81 | 0.24 | 1.32 |
| 6 | Farkas et al. (2015) | TCC | 1 | 2.5 | 1.2 | 1.8 | 0.60 | 1.20 |
| 13 | Zhao et al. (2018) | TCC | NR | NR | NR | NR | 0.10 | 0.86 |

*AP, anterior-posterior coordinate, DV, dorsal-ventral coordinate, ML, medial-lateral coordinate, NR, not reported, TCC, trigeminocervical complex.*

Table 10: Coordinates of thalamic neurons: ranges

| Study.ID | Reference | Recording site | AP min | AP max | ML min | ML max | DV min | DV max |
| --- | --- | --- | --- | --- | --- | --- | --- | --- |
| 10 | Sokolov et al. (2013) | Thalamic nuclei | 2.3 | 3.5 | 2.5 | 3.3 | 4.6 | 5.5 |

*AP, anterior-posterior coordinate, DV, dorsal-ventral coordinate, ML, medial-lateral coordinate.*

Table 11: Coordinates of TCC neurons: averages

| Study.ID | Reference | Recording site | AP average | AP error | ML average | ML error | DV average | DV error |
| --- | --- | --- | --- | --- | --- | --- | --- | --- |
| 3 | Bergerot et al. (2007) | TCC | NR | NR | NR | NR | 0.831 | 0.042 |
| 8 | Oliveira et al. (2016) | TCC | NR | NR | NR | NR | 0.476 | 0.027 |

*AP, anterior-posterior coordinate, DV, dorsal-ventral coordinate, ML, medial-lateral coordinate, NR, not reported, TCC, trigeminocervical complex.*

Histological verification of the recording sites was performed in 4 studies, including two studies with recording in the ventroposteromedial thalamic nucleus.

Table 12: Сharacteristics the studied neuronal populations

| Study.ID | Reference | Recording.site | Recorded.units | Cutaneous.receptive.fields | Modality | Input |
| --- | --- | --- | --- | --- | --- | --- |
| 1 | Akerman and Romero-Reyes (2019) | TCC | neuronal clusters | V1, V2, V3 | WDR | A delta and C-fibres |
| 2 | Andreou and Goadsby (2011) | TCC | single neurons | V1 | WDR | A delta and C-fibres |
| 2 | Andreou and Goadsby (2011) | Thalamic nuclei | single neurons | V1, V2 | WDR, NS | A delta and C-fibres |
| 3 | Bergerot et al. (2007) | TCC | single neurons | V1 | WDR | A delta-fibres |
| 4 | Cumberbatch et al. (1997) | TCC | single neurons | NR | NR | A delta-fibres |
| 5 | Dolgorukova et al. (2020) | TCC | single neurons | V1, V2 | WDR | A delta-fibres |
| 6 | Farkas et al. (2015) | TCC | neuronal clusters | V1 | WDR | A delta and C-fibres |
| 7 | Hoffmann et al. (2019) | TCC | single neurons | V1 | WDR | NR |
| 8 | Oliveira et al. (2016) | TCC | single neurons | V1 | WDR | A delta-fibres |
| 9 | Sokolov et al. (2008) | TCC | single neurons | V1, V2 | NR | A delta and C-fibres |
| 10 | Sokolov et al. (2013) | Thalamic nuclei | single neurons | V1, V2, V3 | WDR | NR |
| 11 | Summ et al. (2021) | TCC | single neurons | NR | WDR | A delta-fibres |
| 12 | Vila-Pueyo et al. (2021) | TCC | neuronal clusters | V1 | NR | A delta and C-fibres |
| 13 | Zhao et al. (2018) | TCC | single neurons | V1 | WDR | A delta and C-fibres |

*Note: The column “input” indicates the types of afferent fibres from which the recorded units received meningeal inputs, NR, not reported, NS, nociceptive-specific units (responsive to noxious mechanical stimulation of cutaneous receptive fields only), TCC, trigeminocervical complex, V1-V3, ophthalmic (V1), maxillary (V2), and mandibular (V3) divisions of the trigeminal nerve, WDR, wide dynamic range units (responsive to both noxious and non-noxious mechanical stimulation of cutaneous receptive fields). Most studies reported additional experiments, which were not eligible for this review. In these cases, the characteristics of the neurons were reported for all studied units.*

Table 13: Baseline values: ongoing activity

| Study.ID | Reference | Recording.site | Modality | Input | Recorded.units | Mean, spikes/s |
| --- | --- | --- | --- | --- | --- | --- |
| 1 | Akerman and Romero-Reyes (2019) | TCC | WDR | A delta and C-fibres | neuronal clusters | 20.4 |
| 5 | Dolgorukova et al. (2020) | TCC | WDR | A delta-fibres | single neurons | 18.1 |
| 6 | Farkas et al. (2015) | TCC | WDR | A delta and C-fibres | neuronal clusters | 23.7 |
| 9 | Sokolov et al. (2008) | TCC | NR | A delta and C-fibres | single neurons | 13.0 |
| 10 | Sokolov et al. (2013) | Thalamic nuclei | WDR | NR | single neurons | 11.2 |

*NR, not reported, TCC, trigeminocervical complex, WDR, wide dynamic range units.*

Table 14: Baseline values: DS-evoked activity

| Study ID | Reference | Recording site | Modality | Input | Recorded units | Mean, spikes/stumulus |
| --- | --- | --- | --- | --- | --- | --- |
| 1 | Akerman and Romero-Reyes (2019) | TCC | WDR | C-fibres | neuronal clusters | 1.6 |
| 1 | Akerman and Romero-Reyes (2019) | TCC | WDR | A delta-fibres | neuronal clusters | 8.2 |
| 5 | Dolgorukova et al. (2020) | TCC | WDR | A delta-fibres | single neurons | 2.5 |
| 6 | Farkas et al. (2015) | TCC | WDR | C-fibres | neuronal clusters | 8.4 |
| 6 | Farkas et al. (2015) | TCC | WDR | A delta-fibres | neuronal clusters | 8.5 |
| 10 | Sokolov et al. (2013) | Thalamic nuclei | WDR | NR | single neurons | 7.2 |

*NR, not reported, TCC, trigeminocervical complex, WDR, wide dynamic range units.*

### 5. Drugs

Table 15: The indication of drugs used in 21 experiments with DS-evoked activity measurement and in 14 experiments with ongoing activity measurement

| Drug.indication | DS-evoked activity | Ongoing activity |
| --- | --- | --- |
| Abortive | 10 | 3 |
| Preventive | 11 | 11 |

Table 16: Studied drugs

| Study.ID | Reference | Drug | Drug.indication | Dose | Vehicle | Control.substance | Route.of.administration |
| --- | --- | --- | --- | --- | --- | --- | --- |
| 1 | Akerman and Romero-Reyes (2019) | Amitriptyline | Preventive | 5 mg/kg | saline | saline | intravenous |
| 1 | Akerman and Romero-Reyes (2019) | Flunarizine | Preventive | 5 mg/kg | 45% HBC | 45% HBC | intravenous |
| 1 | Akerman and Romero-Reyes (2019) | Propranolol | Preventive | 5 mg/kg | saline | saline | intravenous |
| 1 | Akerman and Romero-Reyes (2019) | Valproate | Preventive | 100 mg/kg | saline | saline | intravenous |
| 2 | Andreou and Goadsby (2011) | Topiramate | Preventive | 30 mg/kg | distilled water for injection | distilled water for injection | intravenous |
| 3 | Bergerot et al. (2007) | Naratriptan | Abortive | 5 mg/kg | saline | saline | intravenous |
| 4 | Cumberbatch et al. (1997) | Rizatriptan | Abortive | 0.3, 1 and 3 mg/kg, 10 min apart | saline | saline | intravenous |
| 5 | Dolgorukova et al. (2020) | Metoclopramide | Abortive | 5mg/kg trice, 30 min apart | saline | saline | intravenous |
| 6 | Farkas et al. (2015) | Propranolol | Preventive | 10 mg/kg | saline | saline | subcutaneous |
| 6 | Farkas et al. (2015) | Sumatriptan | Abortive | 1 mg/kg | saline | saline | subcutaneous |
| 6 | Farkas et al. (2015) | Topiramate | Preventive | 30 mg/kg | saline | saline | subcutaneous |
| 7 | Hoffmann et al. (2019) | Magnesium | Abortive | 100 mg/kg | water for injection USP | water for injection USP | intravenous |
| 7 | Hoffmann et al. (2019) | Naratriptan | Abortive | 5 mg/kg | water for injection USP | water for injection USP | intravenous |
| 8 | Oliveira et al. (2016) | Naratriptan | Abortive | 10 mg/kg | sterile water | sterile water | intravenous |
| 9 | Sokolov et al. (2008) | Valproate | Preventive | 200 mg/kg | saline | saline | intravenous |
| 10 | Sokolov et al. (2013) | Valproate | Preventive | 300 mg/kg | NR | saline | intravenous |
| 11 | Summ et al. (2021) | Ibuprofen | Abortive | 30 mg/kg | water for injection | water for injection | intravenous |
| 11 | Summ et al. (2021) | Naproxen | Abortive | 30 mg/kg | water for injection | water for injection | intravenous |
| 12 | Vila-Pueyo et al. (2021) | Lasmiditan | Abortive | 5 mg/kg | saline | saline | intravenous |
| 13 | Zhao et al. (2018) | Propranolol | Preventive | 3 mg/kg | saline | saline | intravenous |

*HBC, 2-hydroxypropyl-beta-cylcodextrin aqueous solution.*

The time of the maximal effect varied between 5 - 90 min after the drug administration.

### 6. Outcomes

The first outcome assessment was performed from 0 to 15 min after treatment and the last outcome assessment was made 30 to 120 min after treatment.

Table 17: The time of the maximal effect and the recording

| Study ID | Experiment ID | Reference | Drug | Outcome | First outcome assessment | Time of the maximal effect | Last outcome assessment |
| --- | --- | --- | --- | --- | --- | --- | --- |
| 1 | 1 | Akerman and Romero-Reyes (2019) | Amitriptyline | Ongoing activity | 15 | 15 | 60 |
| 1 | 1 | Akerman and Romero-Reyes (2019) | Amitriptyline | DS-evoked activity | 15 | 15 | 60 |
| 1 | 2 | Akerman and Romero-Reyes (2019) | Flunarizine | Ongoing activity | 15 | 30 | 60 |
| 1 | 2 | Akerman and Romero-Reyes (2019) | Flunarizine | DS-evoked activity | 15 | 45 | 60 |
| 1 | 3 | Akerman and Romero-Reyes (2019) | Propranolol | Ongoing activity | 15 | 15 | 60 |
| 1 | 3 | Akerman and Romero-Reyes (2019) | Propranolol | DS-evoked activity | 15 | 30 | 60 |
| 1 | 4 | Akerman and Romero-Reyes (2019) | Valproate | Ongoing activity | 15 | 45 | 60 |
| 1 | 4 | Akerman and Romero-Reyes (2019) | Valproate | DS-evoked activity | 15 | 15 | 60 |
| 2 | 5 | Andreou and Goadsby (2011) | Topiramate | DS-evoked activity | 5 | 50 | 90 |
| 2 | 5 | Andreou and Goadsby (2011) | Topiramate | Ongoing activity | 5 | 60 | 90 |
| 2 | 6 | Andreou and Goadsby (2011) | Topiramate | DS-evoked activity | 5 | 40 | 90 |
| 2 | 6 | Andreou and Goadsby (2011) | Topiramate | Ongoing activity | 5 | 70 | 90 |
| 3 | 7 | Bergerot et al. (2007) | Naratriptan | DS-evoked activity | 5 | 30 | 30 |
| 4 | 8 | Cumberbatch et al. (1997) | Rizatriptan | DS-evoked activity | 0 | 20 | 80 |
| 5 | 9 | Dolgorukova et al. (2020) | Metoclopramide | Ongoing activity | 10 | 70 | 90 |
| 5 | 9 | Dolgorukova et al. (2020) | Metoclopramide | DS-evoked activity | 10 | 70 | 90 |
| 6 | 10 | Farkas et al. (2015) | Propranolol | Ongoing activity | 5 | 30 | 30 |
| 6 | 10 | Farkas et al. (2015) | Propranolol | DS-evoked activity | 5 | 30 | 30 |
| 6 | 11 | Farkas et al. (2015) | Sumatriptan | Ongoing activity | 5 | 30 | 30 |
| 6 | 11 | Farkas et al. (2015) | Sumatriptan | DS-evoked activity | 5 | 30 | 30 |
| 6 | 12 | Farkas et al. (2015) | Topiramate | Ongoing activity | 5 | 30 | 30 |
| 6 | 12 | Farkas et al. (2015) | Topiramate | DS-evoked activity | 5 | 25 | 30 |
| 7 | 13 | Hoffmann et al. (2019) | Magnesium | DS-evoked activity | 5 | 30 | 45 |
| 7 | 14 | Hoffmann et al. (2019) | Naratriptan | DS-evoked activity | 5 | 30 | 45 |
| 8 | 15 | Oliveira et al. (2016) | Naratriptan | DS-evoked activity | 0 | 30 | 60 |
| 9 | 16 | Sokolov et al. (2008) | Valproate | DS-evoked activity | 5 | 5 | 45 |
| 9 | 16 | Sokolov et al. (2008) | Valproate | Ongoing activity | 5 | 45 | 45 |
| 10 | 17 | Sokolov et al. (2013) | Valproate | Ongoing activity | 5 | 10 | 90 |
| 10 | 17 | Sokolov et al. (2013) | Valproate | DS-evoked activity | 5 | 5 | 90 |
| 11 | 18 | Summ et al. (2021) | Ibuprofen | DS-evoked activity | 5 | 30 | 45 |
| 11 | 19 | Summ et al. (2021) | Naproxen | DS-evoked activity | 5 | 30 | 45 |
| 12 | 20 | Vila-Pueyo et al. (2021) | Lasmiditan | Ongoing activity | 5 | 90 | 90 |
| 12 | 20 | Vila-Pueyo et al. (2021) | Lasmiditan | DS-evoked activity | 5 | 75 | 90 |
| 13 | 21 | Zhao et al. (2018) | Propranolol | DS-evoked activity | 0 | 60 | 120 |
| 13 | 21 | Zhao et al. (2018) | Propranolol | Ongoing activity | 0 | 60 | 120 |
