## Supplement file 3: Analysis for "Meta-analysis of the effects of clinically-effective therapeutics in the preclinical migraine model as a tool for design optimisation"

Antonina Dolgorukova, Ekaterina Protsenko, Julia Isaeva, Victoria Gagloeva, Elena Verbitskaya, Alexey Y. Sokolov.

The analysis included 13 studies assessing the effects of 13 anti-migraine drugs on the ongoing (8 studies, 14 experiments) and dural stimulation-evoked (DS-evoked, 13 studies, 21 experiments) activity of 290 neurons/neuronal clusters in the electrophysiological model of trigeminovascular nociception (EMTVN).

An individual experiment (defined as a drug tested using a particular route of administration on an animal/neuronal cohort) served as an observational unit in the analyses. We used the absolute difference in means (MD) as an effect size measure and the formulas 5 and 6 in Vesterinen et al. (2014) to calculate the variances of the effect sizes.

In 2 studies (7 experiments), the authors separately analysed A$\delta$ and C-fibre responses (different measures of the evoked activity). We combined these data to obtain a single effect per experiment using the formulas 24 and 25 in Vesterinen et al. (2014).

There were 5 studies producing dependent effect size estimates (multiple experiments per study with a shared control group in 4 cases), thus to calculate the overall effects of the anti-migraine drugs, we fitted a three-level meta-analytic model and computed robust variance estimates.

### 1. Ongoing activity

Table 1: Included experiments

| Study.ID | Exp.ID | Reference | Recording.site | Drug | Route.of.administration |
| --- | --- | --- | --- | --- | --- |
| 1 | 1 | Akerman and Romero-Reyes (2019) | TCC | Amitriptyline | intravenous |
| 1 | 2 | Akerman and Romero-Reyes (2019) | TCC | Flunarizine | intravenous |
| 1 | 3 | Akerman and Romero-Reyes (2019) | TCC | Propranolol | intravenous |
| 1 | 4 | Akerman and Romero-Reyes (2019) | TCC | Valproate | intravenous |
| 2 | 5 | Andreou and Goadsby (2011) | TCC | Topiramate | intravenous |
| 2 | 6 | Andreou and Goadsby (2011) | Thalamic nuclei | Topiramate | intravenous |
| 5 | 9 | Dolgorukova et al. (2020) | TCC | Metoclopramide | intravenous |
| 6 | 10 | Farkas et al. (2015) | TCC | Propranolol | subcutaneous |
| 6 | 11 | Farkas et al. (2015) | TCC | Sumatriptan | subcutaneous |
| 6 | 12 | Farkas et al. (2015) | TCC | Topiramate | subcutaneous |
| 9 | 16 | Sokolov et al. (2008) | TCC | Valproate | intravenous |
| 10 | 17 | Sokolov et al. (2013) | Thalamic nuclei | Valproate | intravenous |
| 12 | 20 | Vila-Pueyo et al. (2021) | TCC | Lasmiditan | intravenous |
| 13 | 21 | Zhao et al. (2018) | TCC | Propranolol | intravenous |

*TCC, trigeminocervical complex.*

#### 1.1. Forest plot: all experiments

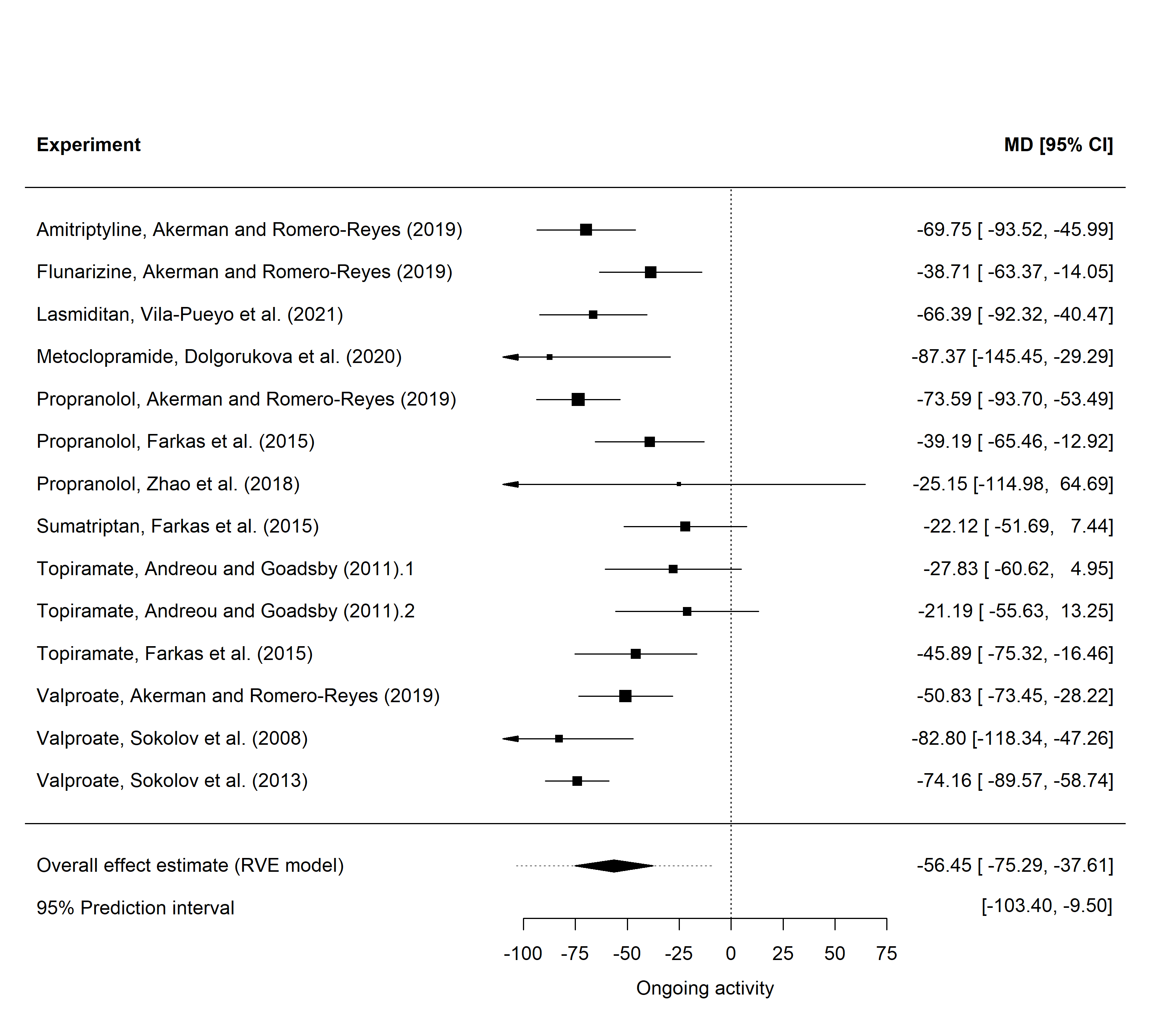

Figure 1: Forest plot of the 14 experiments reporting the drugs’ effects on the ongoing activity from the 8 studies (1-4 experiments per study, mean = 1.8) included in the analysis. The overall effect was calculated using robust variance estimation (RVE). MD, mean difference between treatment and control in %change of neuronal activity from baseline, CI, confidence interval.

The overall I^2^ (which is the sum of between- and within-study heterogeneity) was 64%.

About 54.1% of the total variance was attributed to between-study heterogeneity. The within-study heterogeneity was low (9.8%). The remaining 36% was sampling variance.

**Since overall heterogeneity was moderate, we did not perform subgroup and sensitivity analyses.**

### 2. DS-evoked activity

Table 2: Included experiments

| Study.ID | Exp.ID | Reference | Recording.site | Drug | Route.of.administration |
| --- | --- | --- | --- | --- | --- |
| 1 | 1 | Akerman and Romero-Reyes (2019) | TCC | Amitriptyline | intravenous |
| 1 | 2 | Akerman and Romero-Reyes (2019) | TCC | Flunarizine | intravenous |
| 1 | 3 | Akerman and Romero-Reyes (2019) | TCC | Propranolol | intravenous |
| 1 | 4 | Akerman and Romero-Reyes (2019) | TCC | Valproate | intravenous |
| 2 | 5 | Andreou and Goadsby (2011) | TCC | Topiramate | intravenous |
| 2 | 6 | Andreou and Goadsby (2011) | Thalamic nuclei | Topiramate | intravenous |
| 3 | 7 | Bergerot et al. (2007) | TCC | Naratriptan | intravenous |
| 4 | 8 | Cumberbatch et al. (1997) | TCC | Rizatriptan | intravenous |
| 5 | 9 | Dolgorukova et al. (2020) | TCC | Metoclopramide | intravenous |
| 6 | 10 | Farkas et al. (2015) | TCC | Propranolol | subcutaneous |
| 6 | 11 | Farkas et al. (2015) | TCC | Sumatriptan | subcutaneous |
| 6 | 12 | Farkas et al. (2015) | TCC | Topiramate | subcutaneous |
| 7 | 13 | Hoffmann et al. (2019) | TCC | Magnesium | intravenous |
| 7 | 14 | Hoffmann et al. (2019) | TCC | Naratriptan | intravenous |
| 8 | 15 | Oliveira et al. (2016) | TCC | Naratriptan | intravenous |
| 9 | 16 | Sokolov et al. (2008) | TCC | Valproate | intravenous |
| 10 | 17 | Sokolov et al. (2013) | Thalamic nuclei | Valproate | intravenous |
| 11 | 18 | Summ et al. (2021) | TCC | Ibuprofen | intravenous |
| 11 | 19 | Summ et al. (2021) | TCC | Naproxen | intravenous |
| 12 | 20 | Vila-Pueyo et al. (2021) | TCC | Lasmiditan | intravenous |
| 13 | 21 | Zhao et al. (2018) | TCC | Propranolol | intravenous |

*TCC, trigeminocervical complex.*

#### 2.1. Forest plot: all experiments

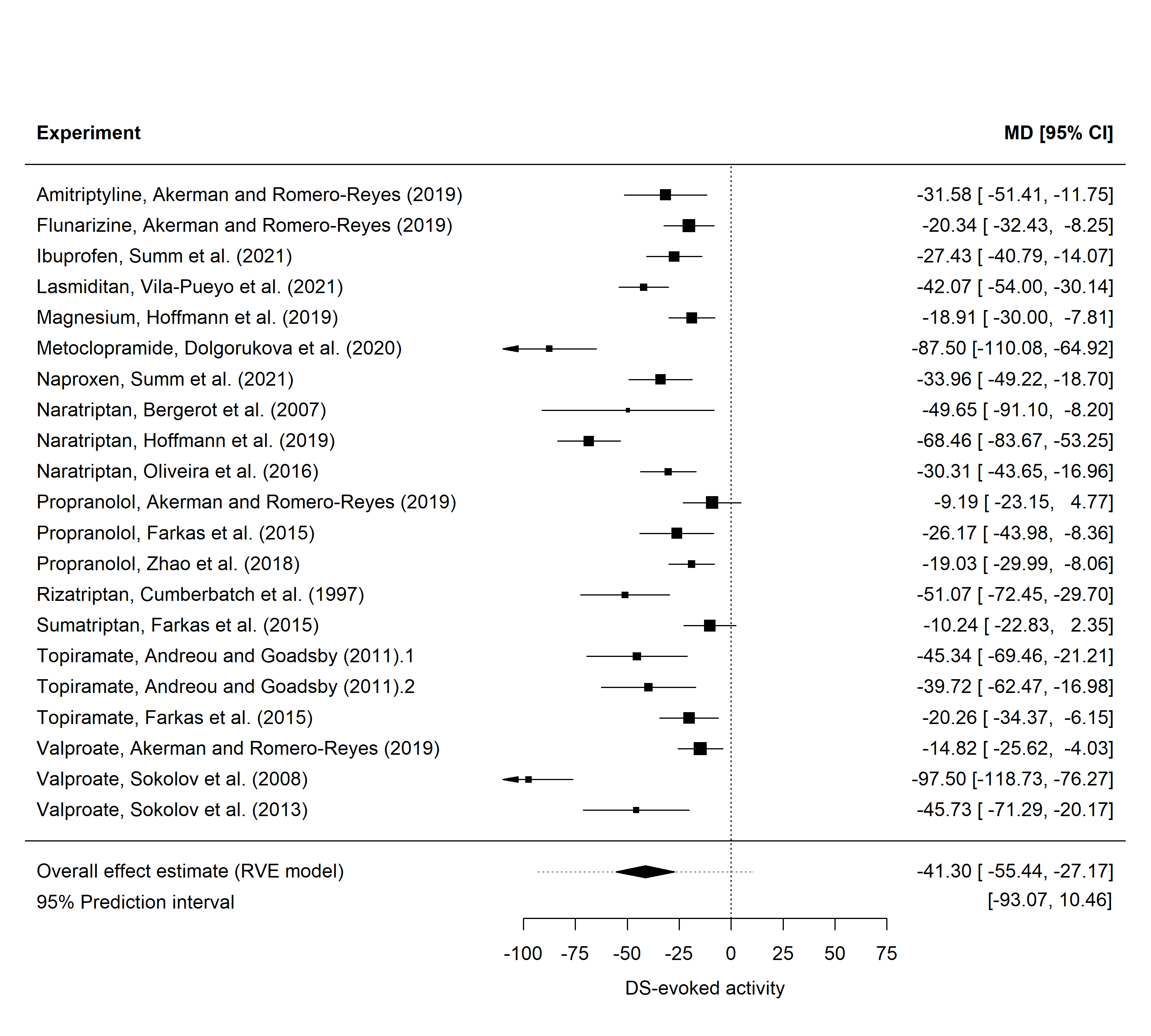

Figure 2: Forest plot of the 21 experiments reporting the drugs’ effects on the DS-evoked activity from the 13 studies (1-4 experiments per study, mean = 1.6) included in the analysis. The overall effect was calculated using robust variance estimation (RVE). MD, mean difference between treatment and control in %change of neuronal activity from baseline, CI, confidence interval.

The overall I^2^ (which is the sum of between- and within-study heterogeneity) was 89.7%.

About 59.2% of the total variance was attributed to between-study heterogeneity. The within-study heterogeneity was low (30.5%). The remaining 10.3% was sampling variance.

#### 2.2. Subgroup analyses

The extent to which methodological features and the reporting of measures to reduce bias explain the observed heterogeneity was assessed in subgroup analyses (meta-regression). In the study protocol we predetermined 7 factors, one of which (sex of the animal) could not be assessed, since all experiments were carried out in male rats.

Table 3: Subgroup analyses (all included studies)

| Moderator | n of experiments | Test for subgroup differences | Overall I^2^ | QE |
| --- | --- | --- | --- | --- |
| Stimulation intensity | 18 | F(1, 9) = 1.29, p = 0.285 | 91.3% | p < 0.001 |
| Anaesthesia induction | 21 | F(2, 10) = 3.41, p = 0.074 | 89.2% | p < 0.001 |
| Anaesthesia maintenance | 21 | F(2, 10) = 1.17, p = 0.348 | 89.7% | p < 0.001 |
| **Gas mixture** | **20** | **F(1, 10) = 11.73, p = 0.006** | **78.2%** | p < 0.001 |
| Drug indication | 21 | F(1, 11) = 0.0014, p = 0.971 | 90.2% | p < 0.001 |
| Time | 21 | F(1, 11) = 0.039, p = 0.847 | 90.2% | p < 0.001 |

*QE, test for residual heterogeneity.*

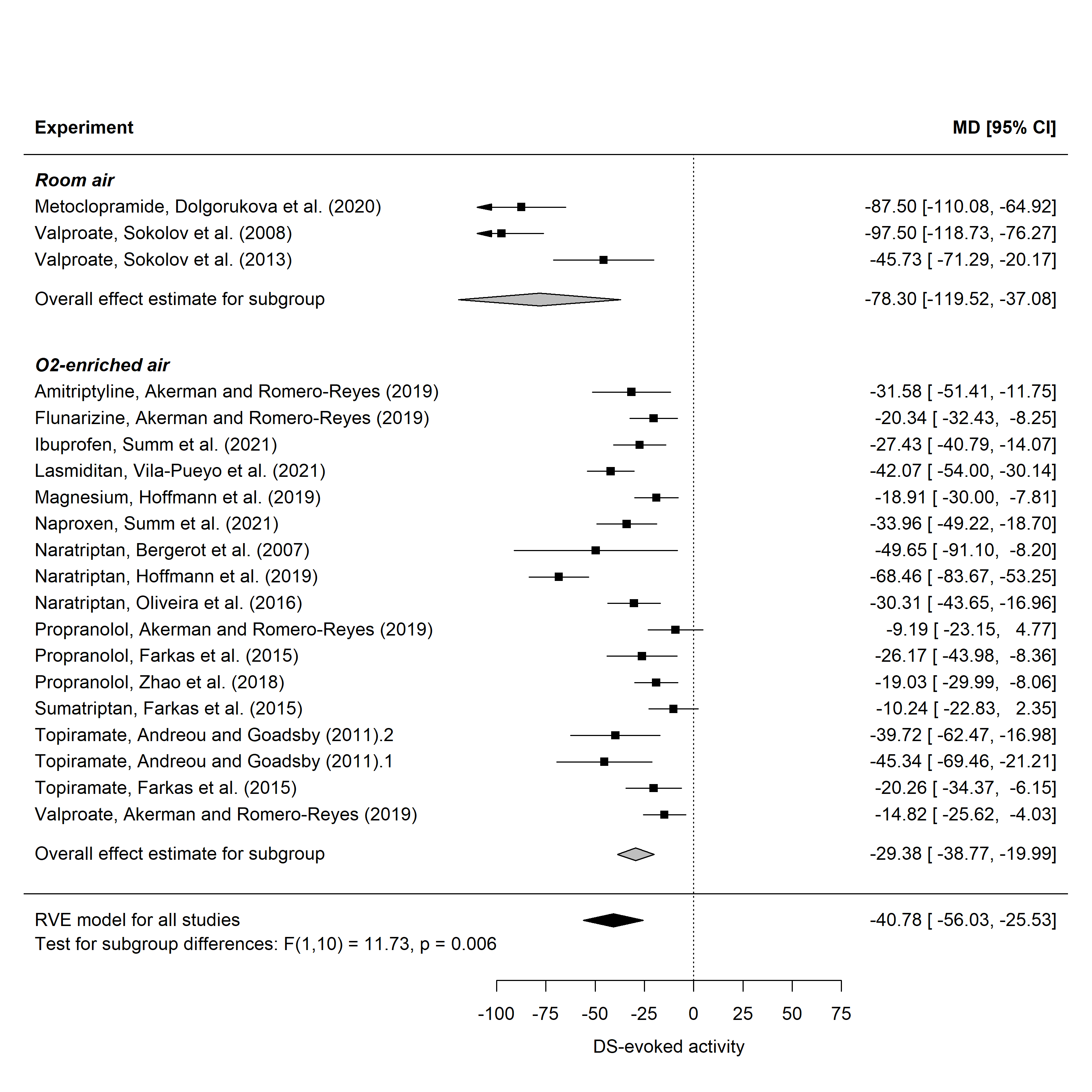

Figure 3: Forest plot of the 20 experiments (12 studies) reporting the drugs’ effects on the DS-evoked activity and the gas mixture used for artificial ventilation by subgroups. MD, mean difference between treatment and control in %change of neuronal activity from baseline, CI, confidence interval.

#### 2.3. Sensitivity analyses

The high heterogeneity in the data set can also be caused by one or more experiments with extreme effect sizes (outliers), while the pooled effect estimate can be highly dependent on a single experiment (influential cases). To detect possible influential outliers we performed sensitivity analyses.

For detecting outliers we calculated standardized (deleted) residuals.

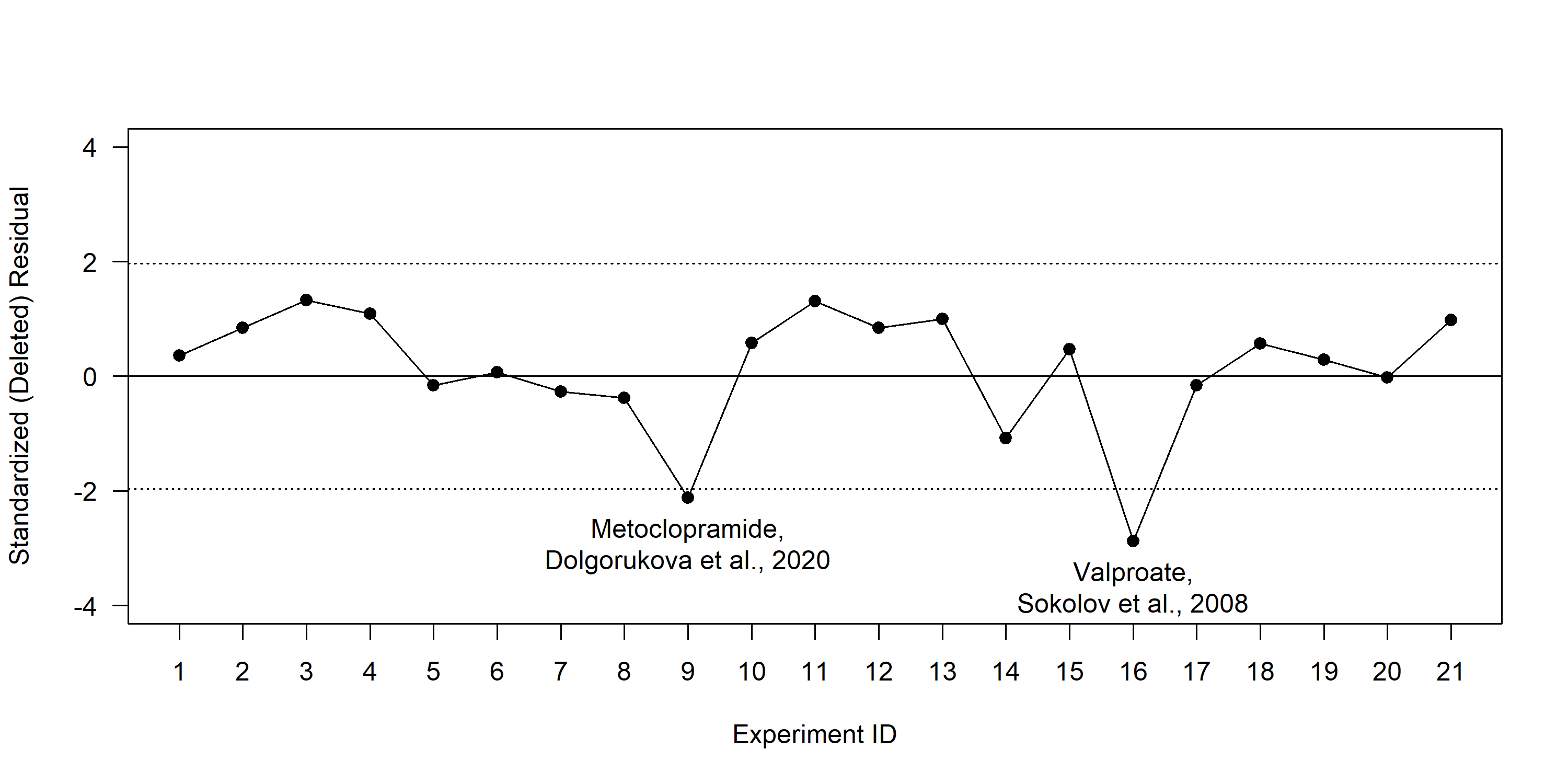

Figure 4: Standardized deleted residuals. The dotted lines represent ±1.96 cut-off. It was assumed that the residuals larger than ±1.96 indicate experiments that does not fit the assumed model, i.e. represent outliers.

We found 2 standardized deleted residuals larger than ±1.96, indicating that the results of these experiments do not fit the assumed model, i.e represent outliers.

For detecting influential cases we constructed the Cook’s distances plot.

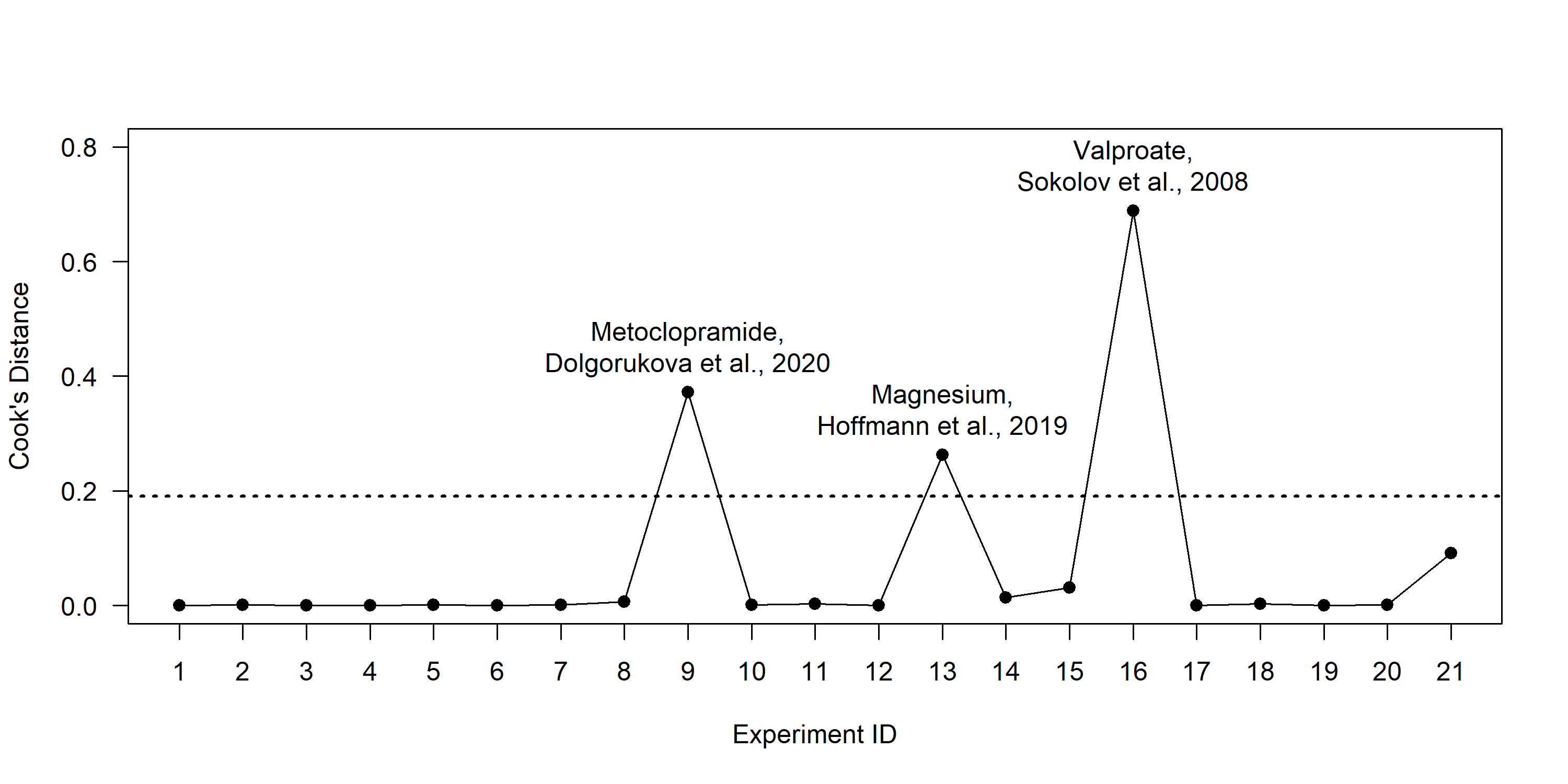

Figure 5: Cook’s distances. The dotted lines represent 4/n, where n is the total number of experiments. It was assumed that Cook’s Distance over that number indicates influential cases.

Thus, out of 21 included experiments, 2 experiments from 2 studies (Dolgorukova et al., 2020, Sokolov et al., 2008) were influential outliers.

The results before and after the removal of the influential outliers are presented below.

Table 4: Results of the sensitivity analysis.

| Analysis | MD | 95%CI | p | Between-study I^2^ | Within-study I^2^ | Overall I^2^ | Sample variance | 95%PI |
| --- | --- | --- | --- | --- | --- | --- | --- | --- |
| All experiments | -41.3 | -55.4 to -27.2 | < 0.001 | 59.2% | 30.5% | 89.7% | 10.3% | -93.1 to 10.5 |
| Outliers removed* | -31.3 | -40.3 to -22.3 | < 0.001 | 20.6% | 56% | 76.5% | 23.5% | -62.9 to 0.2 |

* *The removed influential outliers were metoclopramide tested in Dolgorukova et. al., 2020, and valproate tested in Sokolov et. al., 2008. MD, mean difference between treatment and control in %change of neuronal activity from baseline, CI, confidence interval, PI, prediction interval.*

Since after the exclusion of the experiments that significantly affect the pooled effect estimate overall heterogeneity was still high we rerun subgroup analysis for this data set.

#### 2.4. Subgroup analyses after outliers removal

Table 5: Subgroup analyses (outliers-removed data set)

| Moderator | n of experiments | Test for subgroup differences | Overall I^2^ | QE |
| --- | --- | --- | --- | --- |
| Stimulation intensity | 16 | F(1, 7) = 0.072, p = 0.796 | 69.9% | p < 0.001 |
| **Anaesthesia induction** | **19** | **F(2, 8) = 4.83, p = 0.042** | **74.9%** | p < 0.001 |
| Anaesthesia maintenance | 19 | F(2, 8) = 1.17, p = 0.358 | 77% | p < 0.001 |
| **Gas mixture** | **18** | **F(1, 8) = 14.82, p = 0.005** | **76.8%** | p < 0.001 |
| Drug indication | 19 | F(1, 9) = 2.09, p = 0.182 | 74.3% | p < 0.001 |
| Time | 19 | F(1, 9) = 0.00048, p = 0.983 | 77.5% | p < 0.001 |

*QE, test for residual heterogeneity.*

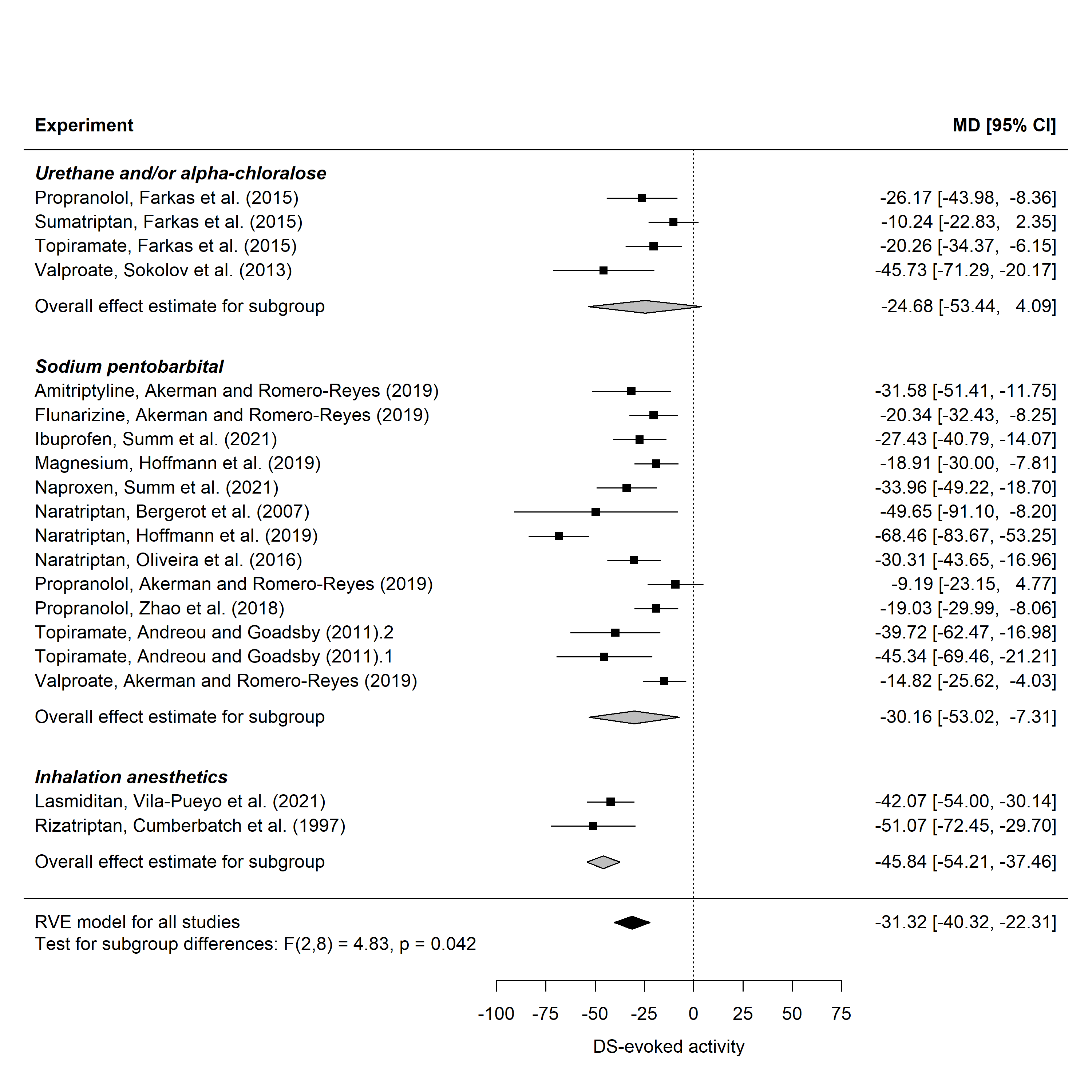

Figure 6: Forest plot of the 19 experiments (11 studies) reporting the drugs’ effects on the DS-evoked activity and the anaesthetic used to initiate anaesthesia by subgroups (outliers-removed data set). MD, mean difference between treatment and control in %change of neuronal activity from baseline, CI, confidence interval.

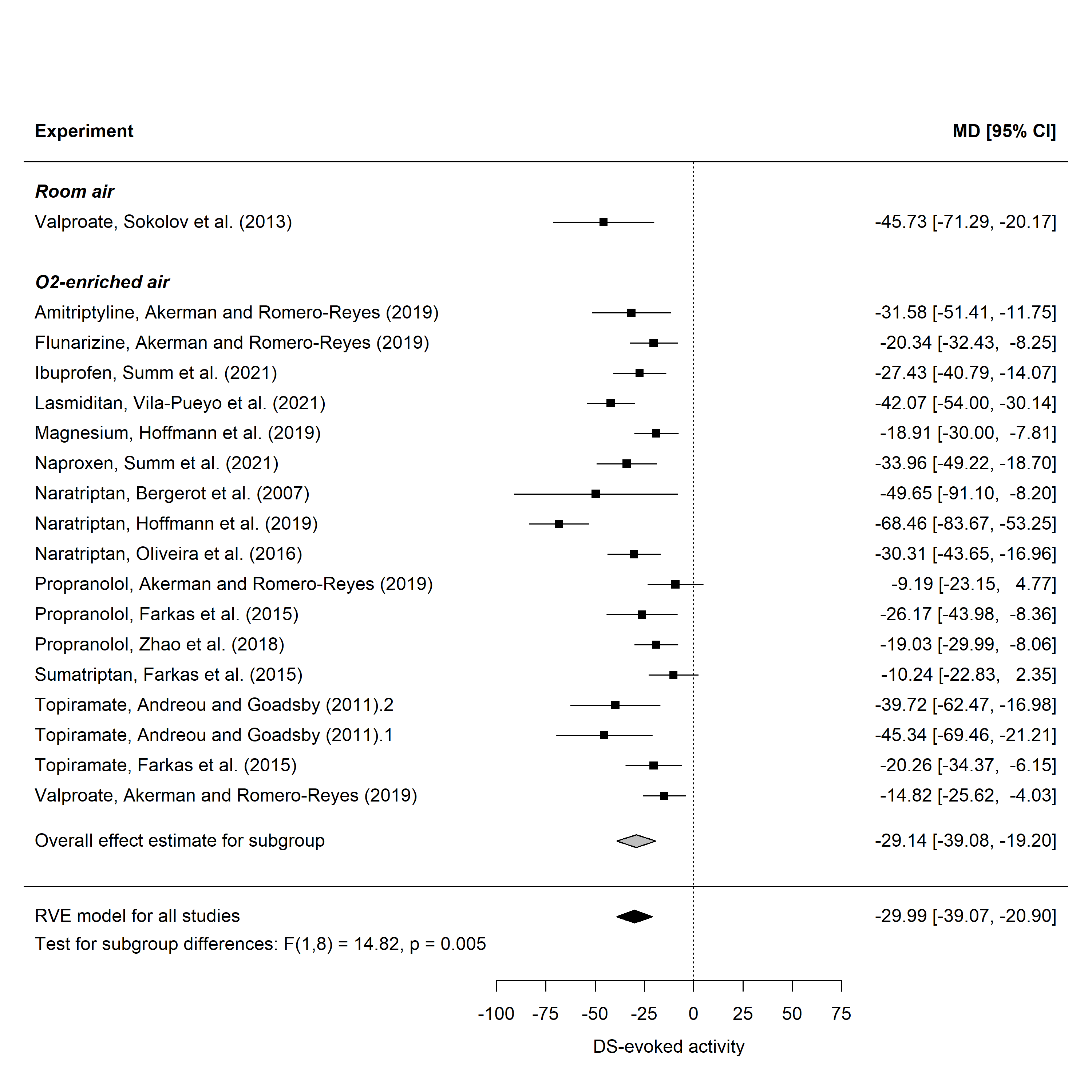

Figure 7: Forest plot of the 18 experiments (11 studies) reporting the drugs’ effects on the DS-evoked activity and the gas mixture used for artificial ventilation by subgroups (outliers-removed data set). MD, mean difference between treatment and control in %change of neuronal activity from baseline, CI, confidence interval.

### 3. Cluster analysis

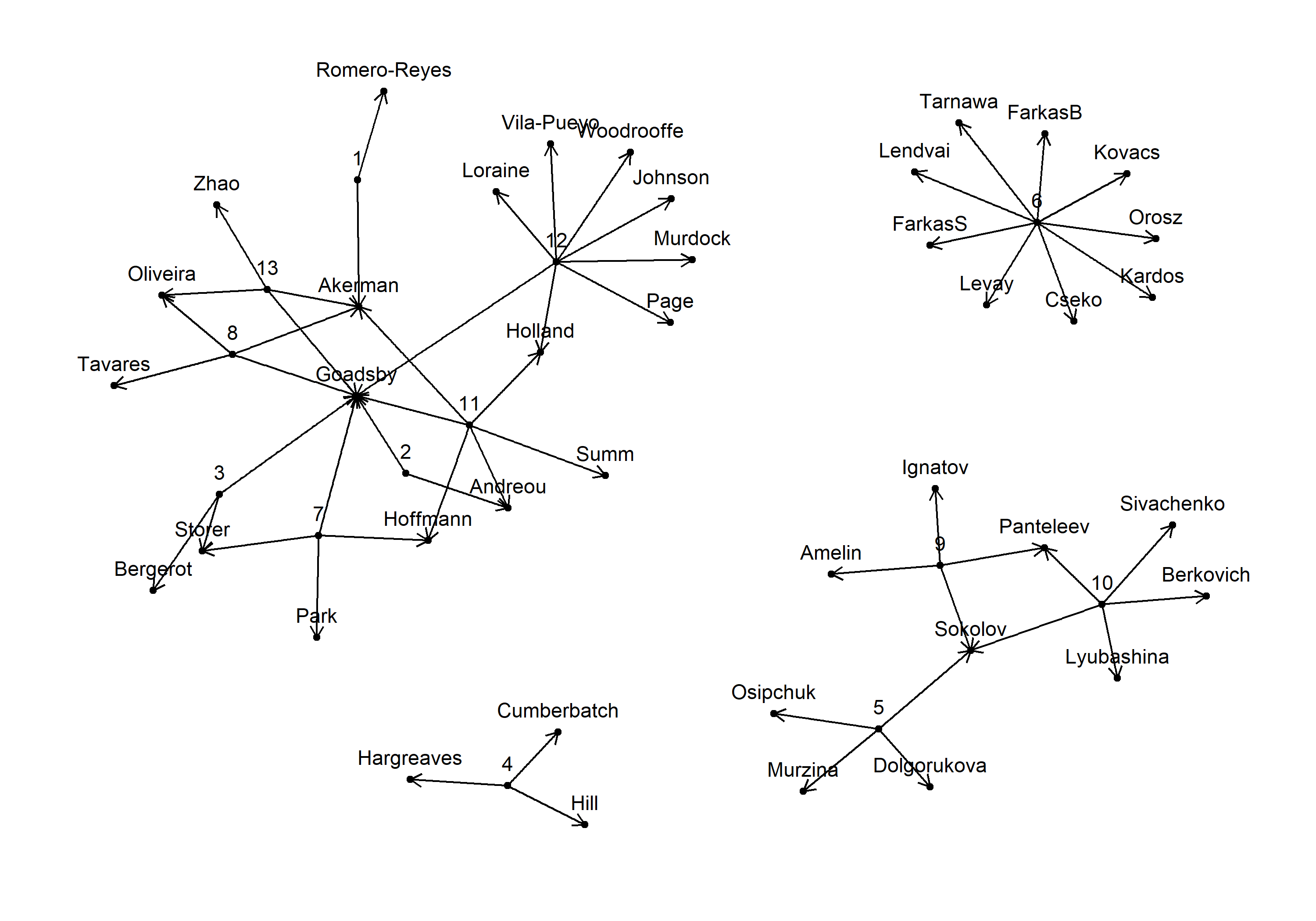

Figure 8: Networks analysis of study’s authors. The numbers represent study ID.

### 4. Risk of Bias

The reporting of some measures to reduce risk of bias and methodological quality criteria may be different for each outcome and have different risk assessment (i.e. attrition or baseline characteristics may be reported for one outcome and not reported for another). However, this was not the case, thus all domains (D) are presented at the study level. The “baseline characteristics” domain assesses reporting the absence of between-group baseline differences and/or accounting for baseline activity fluctuations in the analysis.

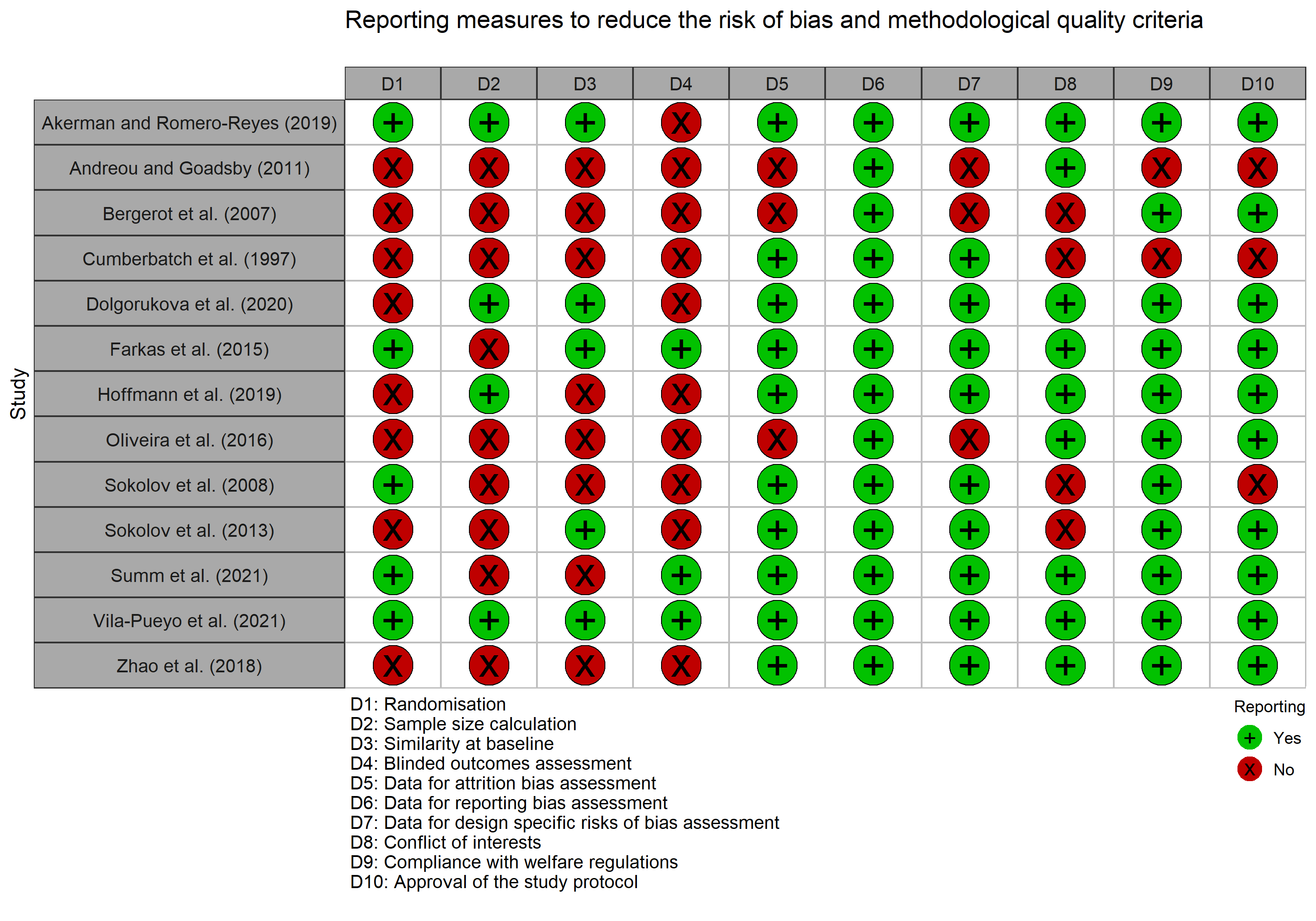

Figure 9: Traffic light plot presenting the reporting measures to reduce the risk of bias and methodological quality criteria in the 13 included studies.

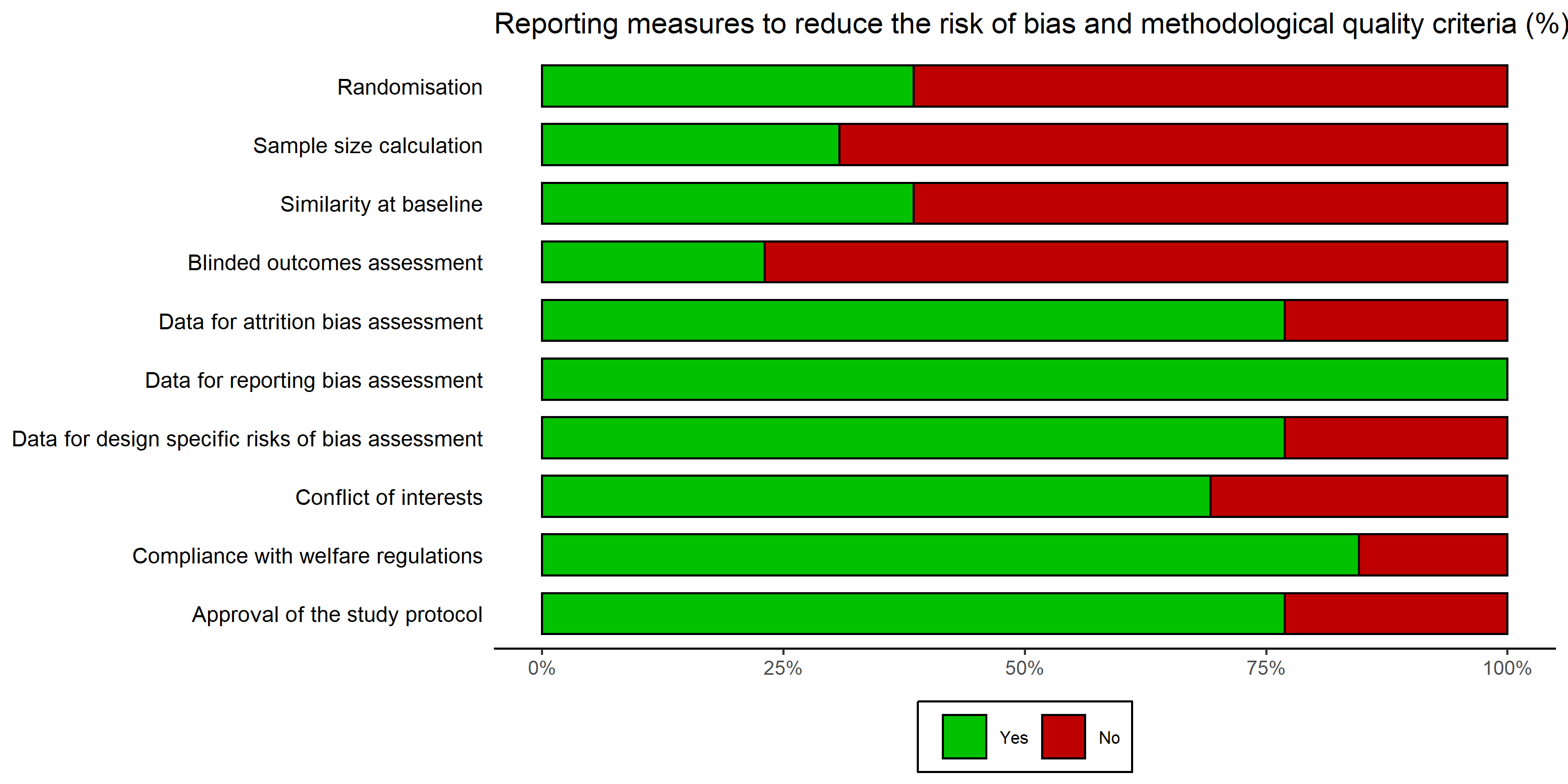

Figure 10: Summary bar plot showing the proportion of studies reporting measures to reduce risk of bias and methodological quality criteria (n = 13).

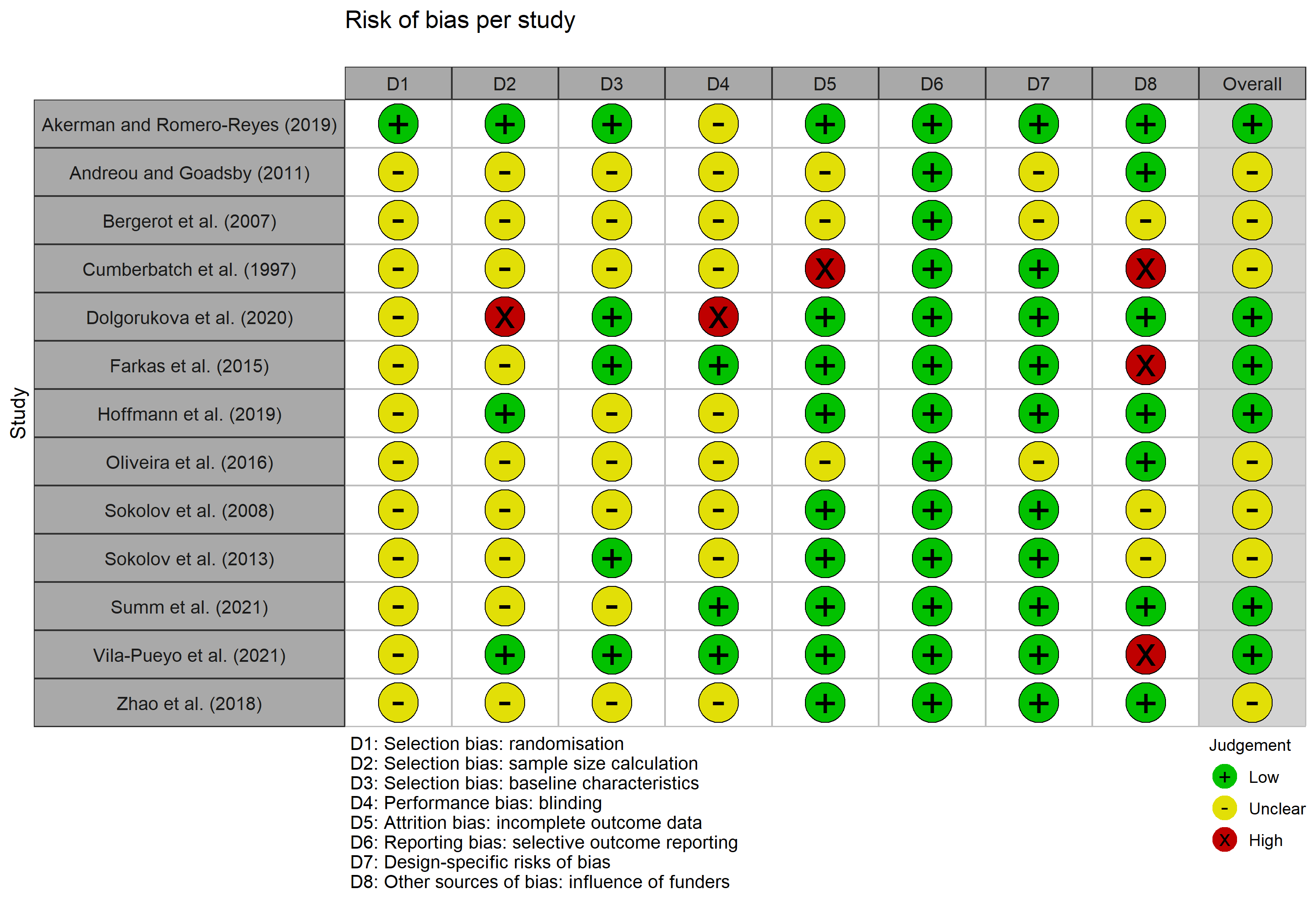

Figure 11: Traffic light plot presenting the risk of bias assigned for each domain for the 13 included studies. The overall risk corresponds to the risk assigned to most domains, in favour of unclear if equal.

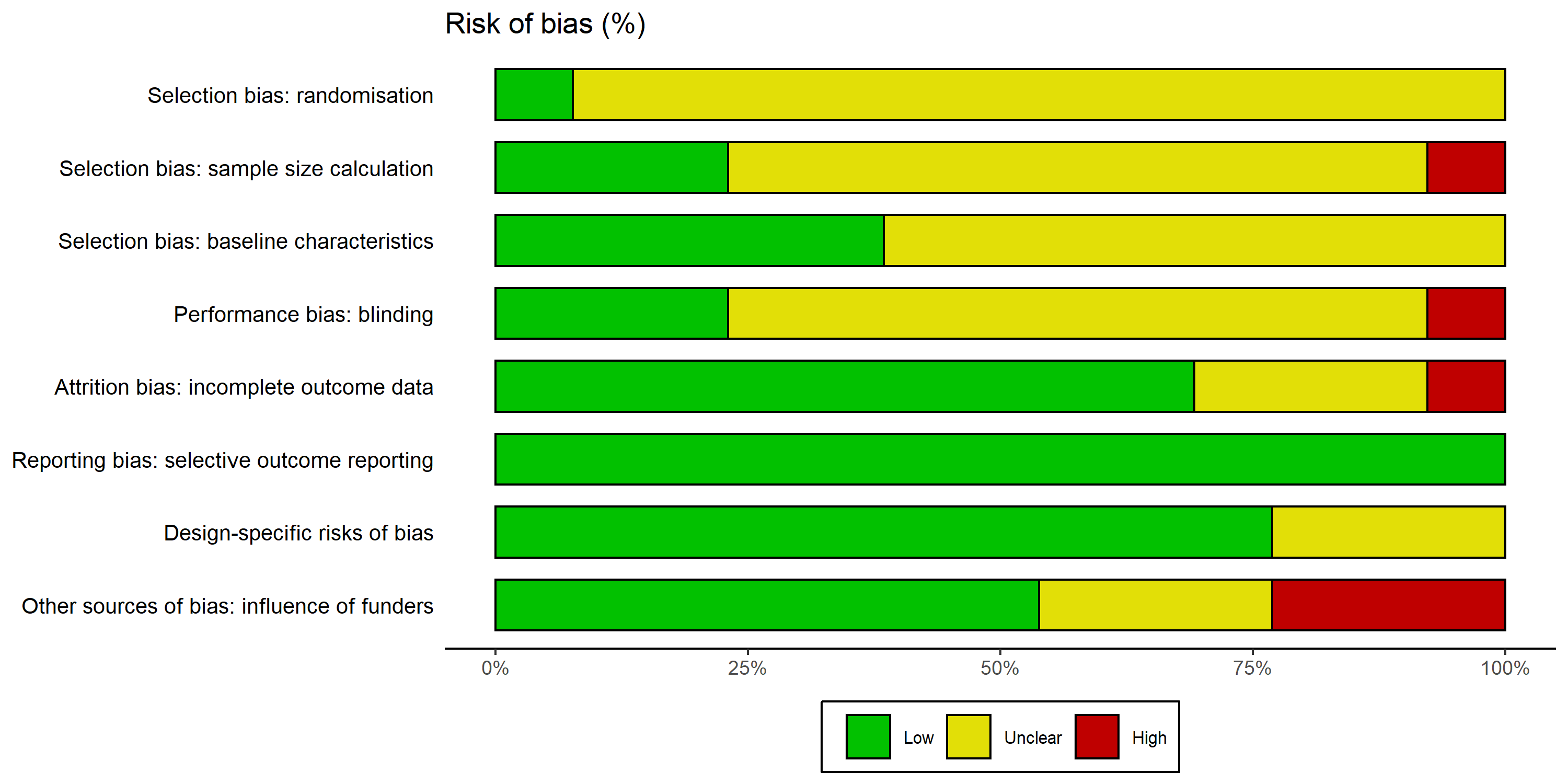

Figure 12: Summary bar plot showing the proportion of studies at low, high, and unclear risk of bias for each criteria (n = 13).

Table 6: Subgroup analyses (all included studies)

| Moderator | n of experiments | Test for subgroup differences | Overall I^2^ | QE |
| --- | --- | --- | --- | --- |
| Randomisation | 21 | F(1, 11) = 0.31, p = 0.591 | 90% | p < 0.001 |
| Sample size calculation | 21 | F(1, 11) = 0.049, p = 0.829 | 90.5% | p < 0.001 |
| Similarity at baseline | 21 | F(1, 11) = 0.16, p = 0.698 | 90.2% | p < 0.001 |
| Blinded outcomes assessment | 21 | F(1, 11) = 2.49, p = 0.143 | 89.6% | p < 0.001 |

*QE, test for residual heterogeneity.*

Table 7: Subgroup analyses (outliers-removed data set)

| Moderator | n of experiments | Test for subgroup differences | Overall I^2^ | QE |
| --- | --- | --- | --- | --- |
| **Randomisation** | **19** | **F(1, 9) = 7.15, p = 0.025** | **72%** | p < 0.001 |
| Sample size calculation | 19 | F(1, 9) = 0.0089, p = 0.927 | 78.1% | p < 0.001 |
| Similarity at baseline | 19 | F(1, 9) = 4.83, p = 0.055 | 72.8% | p < 0.001 |
| Blinded outcomes assessment | 19 | F(1, 9) = 0.45, p = 0.519 | 77.8% | p < 0.001 |

*QE, test for residual heterogeneity.*

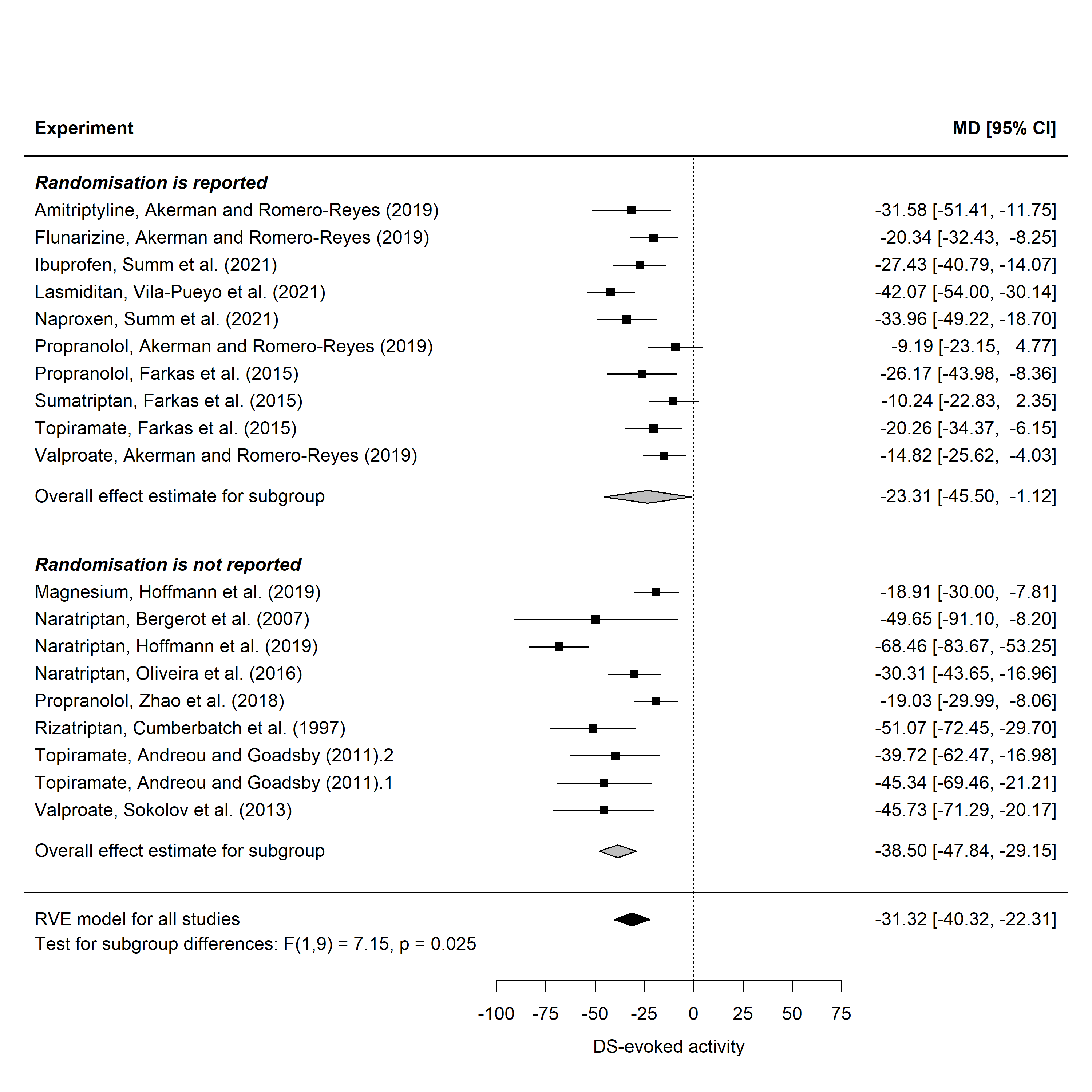

Figure 13: Forest plot of the 19 experiments (11 studies) reporting the drugs’ effects on the DS-evoked activity and grouped by randomisation reporting (outliers-removed data set). MD, mean difference between treatment and control in %change of neuronal activity from baseline, CI, confidence interval.

### 5. Publication Bias

Only the DS-evoked activity data set had a sufficient number of comparisons (≥ 20) to assess a potential publication bias.

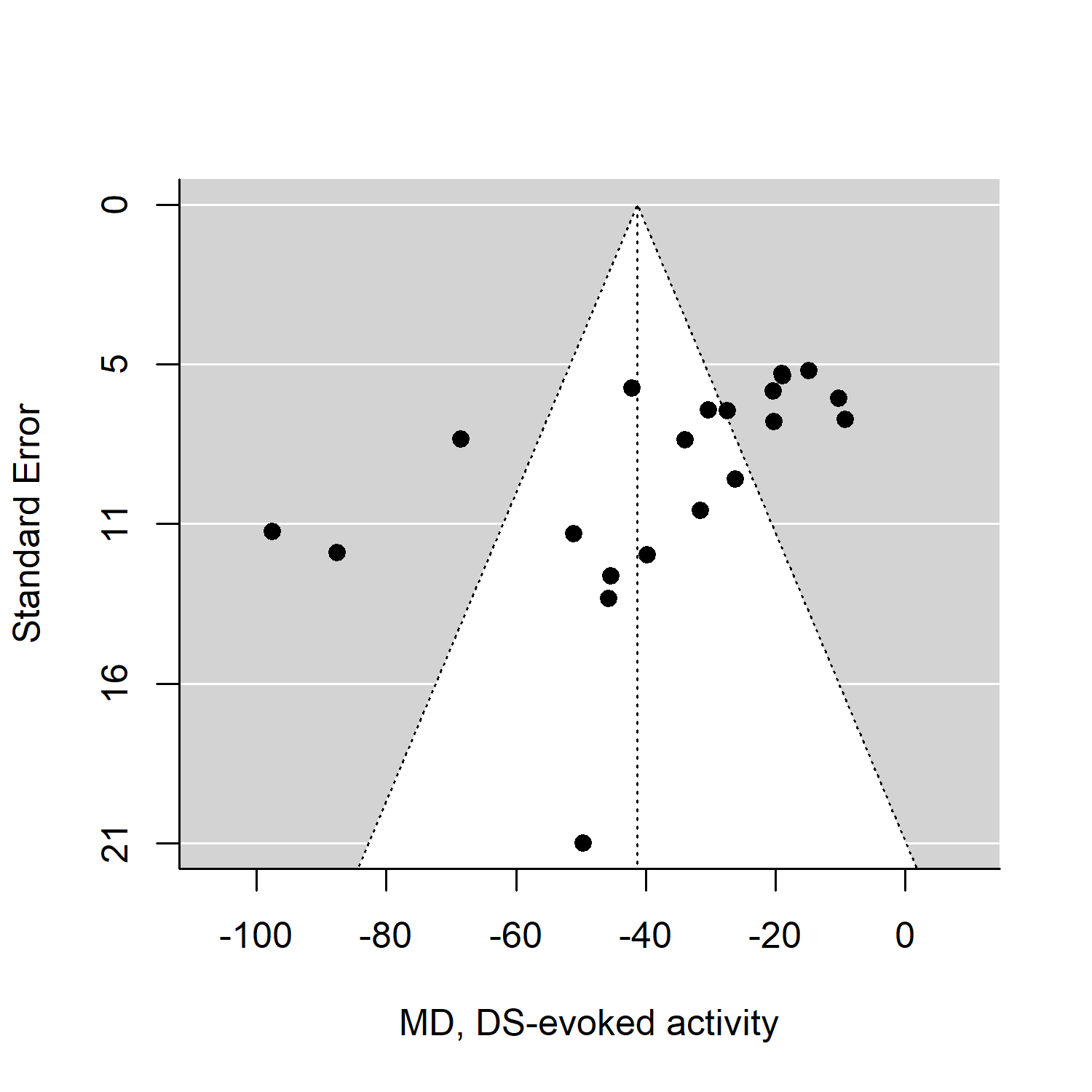

Figure 14: Funnel plot of the 21 experiments reporting the drugs’ effects on the DS-evoked activity extracted from the 13 studies. MD, mean difference between treatment and control in %change of neuronal activity from baseline.

According to Egger’s regression test, there was a significant funnel plot asymmetry (p = 0.033). We could not rerun this analysis without the influential outliers since this dataset include only 19 experiments (≥ 20 needed).

### 6. Sample size calculation

Sample sizes were calculated for the Wilcoxon-Mann-Whitney test at the two-sided alpha = 0.05, 80-100% power, allocation ratio 1:1. In R we used 5% correction to get the same results as

Table 8: The number of animals per group allowing to detect the effect of clinically effective anti-migraine drugs in the EMTVN with 80%-95 power.

| DS-evoked activity Expected MD (rows) / variance (cols) | 13.8% (Small) | 17% (Medium) | 24.5% (Large) | Ongoing activity Expected MD (rows) / variance (cols | 25.2% (Small) | 30.3% (Medium) | 33.6% (Large) |
| --- | --- | --- | --- | --- | --- | --- | --- |
| **Power = 80%** | - | - | - | **Power = 80%** | - | - | - |
| -31.3% (Large) | 5 | 7 | 12 | -56.4% (Large) | 5 | 6 | 8 |
| -25.1% (Medium) | 7 | 9 | 17 | -45.2% (Medium) | 7 | 9 | 11 |
| -15.7% (Small) | 14 | 21 | 42 | -28.2% (Small) | 15 | 21 | 25 |
| **Power = 85%** | - | - | - | **Power = 85%** | - | - | - |
| -31.3% (Large) | 5 | 7 | 13 | -56.4% (Large) | 6 | 7 | 8 |
| -25.1% (Medium) | 7 | 10 | 20 | -45.2% (Medium) | 8 | 10 | 12 |
| -15.7% (Small) | 16 | 24 | 48 | -28.2% (Small) | 17 | 23 | 28 |
| **Power = 90%** | - | - | - | **Power = 90%** | - | - | - |
| -31.3% (Large) | 6 | 8 | 15 | -56.4% (Large) | 6 | 8 | 9 |
| -25.1% (Medium) | 8 | 12 | 23 | -45.2% (Medium) | 9 | 12 | 14 |
| -15.7% (Small) | 19 | 28 | 56 | -28.2% (Small) | 19 | 27 | 33 |
| **Power = 95%** | - | - | - | **Power = 95%** | - | - | - |
| -31.3% (Large) | 7 | 10 | 18 | -56.4% (Large) | 7 | 9 | 11 |
| -25.1% (Medium) | 10 | 14 | 28 | -45.2% (Medium) | 10 | 14 | 17 |
| -15.7% (Small) | 23 | 34 | 69 | -28.2% (Small) | 23 | 33 | 40 |

*MD, mean difference between treatment and control in %change of neuronal activity from baseline.*

*Large, medium, and small expected mean difference (MD, in rows) represents 100%, 80%, and 50% of the overall anti-migraine drug effect estimated with meta-analysis of controlled studies, respectively. Large, medium, and small expected variance values (in columns) are based on cut-off thresholds of pooled standard deviations (SD) that had 80%, 50%, and 20% percent of experiments included in the meta-analysis. For example, 80% of experiments had pooled SD at or below 24.5% and 33.6% for DS-evoked and ongoing activity, respectively. The power analysis for DS-evoked activity was made based on the outlier-removed data set.*
